# Diversification, disparification, and hybridization in the desert shrubs *Encelia*

**DOI:** 10.1101/2020.07.31.230938

**Authors:** Sonal Singhal, Adam B. Roddy, Christopher DiVittorio, Ary Sanchez-Amaya, Claudia L. Henriquez, Craig R. Brodersen, Shannon Fehlberg, Felipe Zapata

**Affiliations:** Department of Biology, CSU Dominguez Hills, 1000 E Victoria Street, Carson CA 90747; School of the Environment, Yale University, New Haven, CT 06511; Institute of Environment, Florida International University, Miami, FL, 33133; University of California Institute for México and the United States, 3324 Olmsted Hall, University of California, Riverside, CA 92521; Pinecrest Research Corporation, 5627 Telegraph Avenue, Suite 420, Oakland, CA 94609; Department of Ecology and Evolutionary Biology, University of California, Los Angeles, 612 Charles E. Young Dr. South, CA 90095; Research, Conservation, and Collections, Desert Botanical Garden, 1201 N Galvin Parkway, Phoenix, AZ 85008

**Keywords:** Aridity, deserts, *Encelia*, hybridization, phylogenomics, trait evolution

## Abstract

- There are multiple hypotheses for the spectacular plant diversity found in deserts. We explore how different factors, including the roles of ecological opportunity and selection, promote diversification and disparification in *Encelia*, a lineage of woody plants in the deserts of the Americas.
- Using a nearly complete species-level phylogeny along with a broad set of phenotypic traits, we estimate divergence times and diversification rates, identify instances of hybridization, quantify trait disparity, and assess phenotypic divergence across environmental gradients.
- We show that *Encelia* originated and diversified recently (mid-Pleistocene) and rapidly, with rates comparable to notable adaptive radiations in plants. *Encelia* probably originated in the hot deserts of North America, with subsequent diversification across steep environmental gradients. We uncover multiple instances of gene flow between species. The radiation of *Encelia* is characterized by fast rates of phenotypic evolution, trait lability, and extreme disparity across environments and between species-pairs with overlapping geographic ranges.
- *Encelia* exemplifies how interspecific gene flow in combination with high trait lability can enable exceptionally fast diversification and disparification across steep environmental gradients.

## Introduction

Despite the seemingly barren landscape of arid habitats, desert ecosystems harbor some of the most spectacular plant evolutionary radiations (Klak *et al.*, 2004; Hernandez-Hernandez *et al.*, 2011). Why and how aridity has promoted the diversification of plant species and phenotypes has puzzled generations of ecologists and evolutionary biologists alike (Stebbins, 1952; Axelrod, 1972), resulting in several hypotheses. First, arid habitats worldwide began to form and expand only in the last 10 my (Arakaki *et al.*, 2011), a relatively short geological time frame during which new regions of niche space became available. Additionally, deserts have been dynamic through space and time, with diverse orogenies, glacial cycles, marine incursions, and volcanic eruptions that likely caused highly variable selection regimes, multiple cycles of migration-isolation, and eventually colonization and diversification in new habitats (Thompson & Anderson, 2000; Riddle *et al.*, 2000; Oskin & Stock, 2003; Conly *et al.*, 2005). Second, in deserts, substantial topographic, edaphic, climatic, and ecological heterogeneity results in a diversity of habitats, across which species can persist and diversify (Ellis *et al.*, 2006; Sosa *et al.*, 2020). Third, the environmental factors that characterize arid ecosystems typically represent extreme conditions for plant functioning and survival, including, but not limited to, drought-stress, high UV radiation, high temperature, and high salinity (Sandquist, 2014). These extreme conditions often occur over narrow geographic regions. When these multiple stressors converge, multiple, functionally equivalent solutions to the same challenge can evolve. This can lead to phenotypic disparification in otherwise seemingly homogeneous environments (Niklas, 1994).

The confluence of these geological, environmental, and ecological factors in arid ecosystems are likely crucial in spurring the radiation of resident plant lineages (Hernández-Hernández *et al.*, 2014; Said Gutiérrez-Ortega *et al.*, 2018), and the multiplicity of strong selective agents that occur in arid ecosystems may be responsible for the remarkable morphological and physiological diversity that have evolved among desert plants. Desert lineages, therefore, provide ideal case studies of the role of selection in plant radiations. Yet, integrative studies that link the evolutionary history of a lineage with patterns of phenotypic, ecological, climatic, and environmental variation are lacking, and thus our understanding of the processes of diversification of plants in arid regions remains elusive. Here, we use an integrative approach to document the evolutionary radiation of shrubs in the genus *Encelia* (Asteraceae), which are widespread throughout the deserts of the Americas, and showcase how multiple factors – abiotic, biogeographic, phenotypic, and population dynamics – interact to produce high diversification within a clade.

Most species of *Encelia* are distributed in the arid lands of southwestern North America, the dry lands of Chile, Peru, and Argentina, and the (arid) Galapagos Islands (Clark, 1998). These plants inhabit various types of deserts, including inland deserts, coastal dunes, as well as high and low deserts; *E. actoni* even passes the frost line in the Sierra Nevada mountain range of California. Given the widespread distribution of *Encelia*, it is plausible that the dynamic geologic and climatic history of the arid habitats has provided multiple opportunities for lineage separation and diversification (Spotila *et al.*, 1998; Dolby *et al.*, 2015). Indeed, previous work has shown that range fragmentation and expansion associated with climatic changes during the Pleistocene have influenced the spatial distribution of genetic diversity in the widespread *Encelia farinosa* (Fehlberg & Ranker, 2009; Fehlberg & Fehlberg, 2017). However, the influence of biogeographic, ecological, or other abiotic forces on the radiation of the genus *Encelia* is unknown.

Commonly referred to as brittlebushes, *Encelia* species display remarkable eco-phenotypic diversity, and their phenotypic traits are strongly associated with habitat differentiation (Ehleringer & Clark, 1988; Clark, 1998). The species range from small to medium-size shrubs (0.2 - 1.5 m in height) but exhibit substantial variation in overall plant architecture. Leaf morphology is exceptionally diverse in *Encelia*. The leaves are always simple and spirally arranged but vary extensively in shape, margin, size, and indumentum. Classic studies in plant ecophysiology have shown fitness tradeoffs between leaf morphological traits and physiological functions that are associated with fine-scale habitat differentiation (Ehleringer *et al.*, 1981; Ehleringer, 1988). In contrast to vegetative structures, inflorescence morphology, floret morphology, and flowering phenology in *Encelia* do not display substantial diversity, consistent with Asteraceae more generally. Thus. much of the phenotypic diversification in *Encelia* occurs among vegetative structures, although the overall tempo and mode of phenotypic evolution – including the rate of trait evolution and correlation in evolution across traits – in *Encelia* is not understood.

As currently circumscribed, *Encelia* includes 15 species and 5 subspecies (Clark, 1998). Most species are allo- or parapatric, but contact zones where natural hybrids form are common throughout the geographic range of *Encelia* (Clark, 1998). Natural hybridization is rampant in *Encelia* (Kyhos, 1967; Kyhos et al., 1981), and some species are hypothesized to result from hybrid speciation (Allan *et al.*, 1997). Nevertheless, all species seemingly maintain their phenotypic cohesion and independence. Though divergent selection can maintain species despite widespread hybridization (DiVittorio *et al.*, 2020), whether interspecific gene flow increases genetic diversity through hybridization or introgression among *Encelia* species and thus enables lineages to take advantage of new ecological opportunities is not known.

Understanding the evolution of the unique ecology, phenotypic diversity, and the role of gene flow in the radiation of *Encelia* requires a well-resolved phylogeny of the group. Variation in morphology, secondary chemistry, and sequence data show that *Encelia* is monophyletic and most closely related to *Gerea* and *Enceliopsis*, however the relationships among the species in the genus have been difficult to resolve (Clark, 1998; Fehlberg & Ranker, 2007). Here we present a broadly sampled phylogenetic analysis of *Encelia* using RAD seq from 12 *Encelia* species and 2 outgroup species. Using this phylogeny, we address four questions: a) what is the evolutionary history of *Encelia*? b) what is the tempo and mode of diversification and trait disparification in *Encelia*? c) what are the main drivers of diversification and trait disparification in *Encelia*? and d) what is the role of interspecific gene flow in this evolutionary radiation?

## Methods

### Sampling

We sampled 77 individuals from 12 of the 15 recognized species in *Encelia* (Fig. 1; Table S1). Where possible, we collected individuals across the range. For all species, we collected leaf and seed material during field seasons ranging from 2009 - 2016. Species were grown in a common garden at the University of California Agricultural Operations Station in Riverside, CA; further details are available in the Supplemental Methods. Phenotypic measurements and tissues for genetic analysis were taken from adult individuals in the garden for all species but *Encelia ravenii* and *E. resinifera*, for which we used field-collected adult leaves.

**Figure 1:**
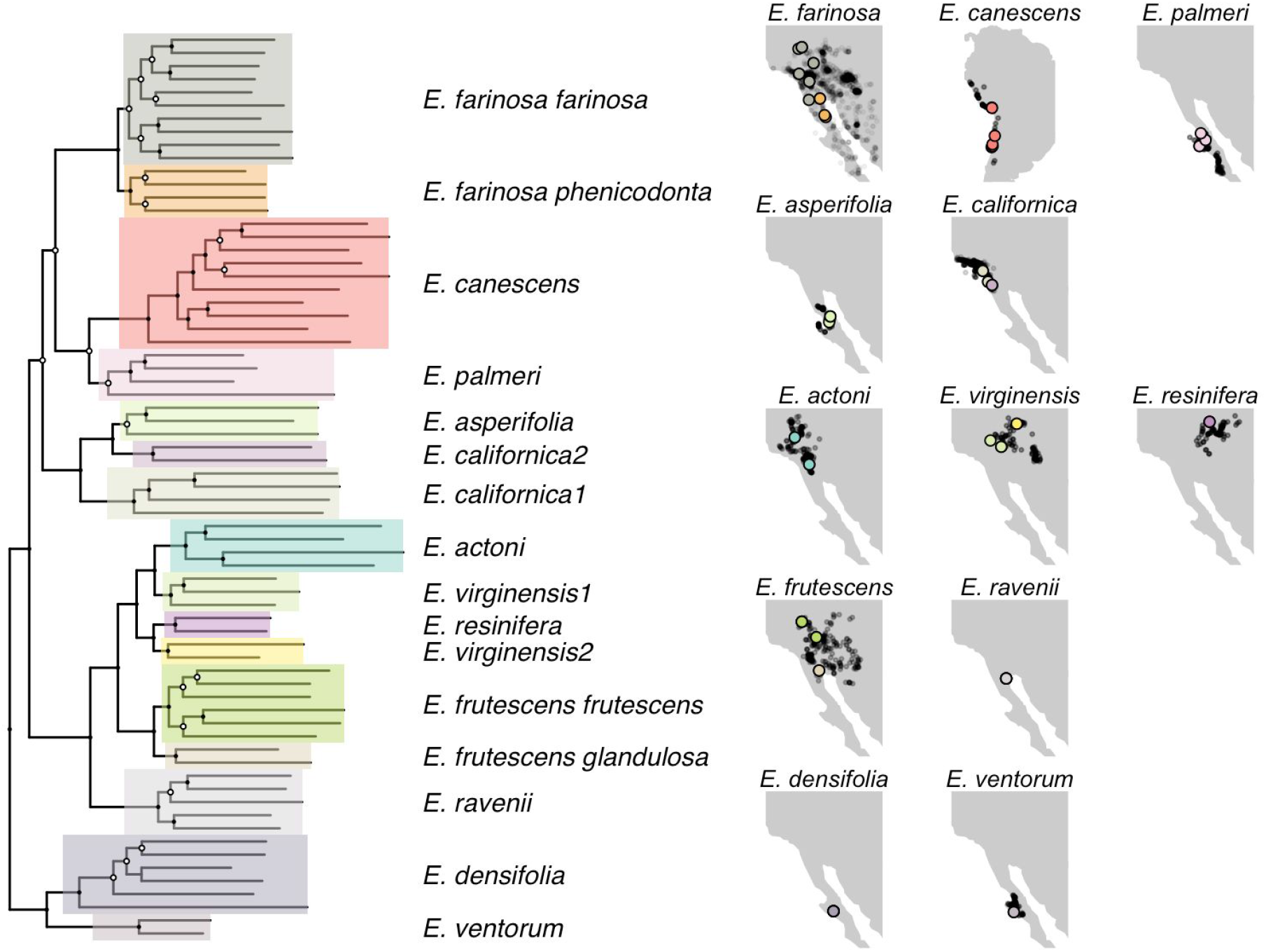
Individual-level phylogeny of *Encelia*, inferred using RAxML on a concatenated alignment of 31K loci. Shaded boxes demarcate clades; nodes with bootstrap <95% are indicated in white. Maps show distribution of nominal species based on GBIF data as light gray points. Large filled points indicate sampling locations for individuals used in this study, colored by lineage identity.

### Genetic data collection

We collected genetic data using double-digest restriction-aided (ddRAD) sequencing. We first extracted DNA from silica-dried adult leaves and then prepared doubly-barcoded ddRAD libraries (Peterson *et al.*, 2012) using *PstI* and *MspI* and size-selecting fragments from 250 - 700 bp. All libraries were pooled and sequenced across one lane of 100 PE sequencing on the Illumina HiSeq 4000 Sequencing Platform.

To process and analyze these data, we wrote a pipeline that generates both pseudo-reference genomes per lineage and variant call sets across individuals within a lineage. This approach generates a common reference index across lineages, making it easier to identify homologous loci and variants. Full details are in the Supplemental Information. First, we cleaned and assembled reads using Trimmomatic v36 (Bolger *et al.*, 2014), PEAR v0.9.8 (Zhang *et al.*, 2014), and Velvet v1.2.10 (Zerbino & Birney, 2008). Second, we determined lineage identity per individual. Although all sampled individuals were identified to nominal species, species boundaries have not been well-tested in *Encelia*. To do so, we used VSEARCH v2.9.1 (Rognes *et al.*, 2016) to identify homologous loci using a 70% identity cutoff. We then aligned loci using mafft v7.9.34 (Katoh *et al.*, 2009), concatenated these loci, and used RAxML v8.2.11 to generate a phylogeny from the 11.6K loci, 1.5 Mb concatenated alignment (Stamatakis, 2014). By comparing clade identity to nominal species identity, we determined likely lineage identities for each individual (Table S1). Finally, after determining lineage identities, we generated a pseudo-reference genome per lineage. To do so, we used an iterative reference-based approach (Sarver *et al.*, 2017). Across all individuals, we selected the longest locus in each homolog group. This unique set of 244K loci served as our starting pseudo-reference genome. Per lineage, we mapped reads to the starting pseudo-reference genome using bwa v0.7.17 (Li, 2013), called variants using samtools v1.5 (Li *et al.*, 2009), and then mutated the current pseudo-reference genome to incorporate any variants at ≥50% allele frequency. This was repeated three additional times to result in a final pseudo-reference genome specific to the lineage. We generated the final variant set per lineage by using bwa to align reads and then calling genotypes using samtools. We retained all sites with quality scores >20 and all genotypes with depth ≥10x.

### Genetic data analyses

#### Phylogenetic inference

We inferred individual-level and lineage-level phylogenies using concatenated and coalescent-based models; full details are available in the Supplemental Information. For an individual-level phylogeny, we used RAxML to infer a phylogeny and 100 bootstrap replicates from a 31K loci, 3.9 Mb concatenated alignment. We used SVDquartets based on 2.8K SNPs to infer a coalescent-based phylogeny with 100 bootstrap replicates (Chifman & Kubatko, 2014). Bootstrap estimates of nodal support are often inflated, particularly for phylogenomic datasets (Cummings *et al.*, 2003). Accordingly, we calculated gene concordance factors (gCF) and site concordance factors (sCF) using IQ-TREE v1.6.4 (Minh *et al.*, 2020), both of which better measure conflict across loci and sites.

For a lineage-level phylogeny, we used two coalescent-based approaches. First, we used ASTRAL-III (Zhang *et al.*, 2018) based on 29K gene trees inferred using RAxML. Second, we used SVDquartets with the same SNP dataset used for the individual-level SVDquartets phylogeny (see above). Neither ASTRAL-III nor SVDquartets provides terminal branch lengths. Accordingly, we used RAxML to estimate branch lengths based on a concatenated alignment on a constrained topology. To infer a time-calibrated phylogeny, we used an external calibration from a comprehensive angiosperm phylogeny that estimated the crown age of *Encelia* as 1.36 million years (Myr) (Magallón *et al.*, 2015; Smith & Brown, 2018). This aligns with previous divergence dating based on population genomic data that inferred the crown of *Encelia* as 1.05 Myr (Singhal, unpublished). We used this root age to infer a chronogram using the ‘chronos’ function in the R package ‘ape’ with lambda of 0.01. We used this time-calibrated phylogeny in all comparative analyses.

#### Introgression

Given previous analyses and field studies have suggested hybridization is common in *Encelia* (Clark & Allan, 1997; Allan *et al.*, 1997), we used two complementary approaches to identify likely instances of historical and current introgression. First, we inferred phylogenetic networks using SNAQ v0.9.0 (Solís-Lemus & Ané, 2016). As input, we provided gene trees, removing any gene trees with >50% missing data. We then ran SNAQ for 0 to 5 reticulate edges, for three independent replicates each, using the ASTRAL tree as the starting topology. Second, we calculated the D-statistic across lineages, which measures when topological variance is in excess of what would be predicted under incomplete lineage sorting (Durand *et al.*, 2011). Using all possible species triads based on the ASTRAL topology with *Enceliopsis covelli* as the outgroup, we calculated an allele frequency based D-statistic. We then calculated significance of the D-statistic by conducting 1000 bootstraps and calculating the Z-score (Eaton & Ree, 2013). For a given species pair, we report the D-statistic calculated using the nearest neighbor as the third lineage. If this still resulted in multiple comparisons, we conservatively report the D-statistic with the smallest Z-score (Malinsky *et al.*, 2018).

#### Population genetics

Given that *Encelia* lineages likely radiated rapidly, genetic divergence should be similar across species comparisons. To test this prediction, we calculated *D*_*xy*_ and *F*_*ST*_ (Nei, 1978; Reich *et al.*, 2009) across all pairwise lineages.

### Trait data collection

We collected nine morphological and physiological traits (summarized in Table S2) across species to determine the extent and nature of phenotypic variation. In addition, we used microCT imaging to characterize the fine-scale external and internal morphology of leaves as well as to calculate trichome density and stem xylem vessel diameter and area. Below we briefly describe how these data were collected; full details are available in the Supplemental Information.

#### Leaf area, shape, and color

To analyze leaf area, shape, and color, we collected and photographed three to five adult leaves per individual growing in the common garden. We analyzed leaf images using ImageJ (Rasband & Others, 1997) and summarized leaf measurements using a scaled and centered principal component analysis. The first two axes summarize 42% and 24% of the variation respectively; PC1 largely loads on size-related measurements and PC2 largely reflects the roundness of the leaf. To measure color, we used the white-balanced leaf images in Adobe Photoshop and measured the arithmetic mean of all pixels of the largest circumscribed rectangle possible within the center of the leaf. Leaf mass was measured on dry leaves and used to calculate leaf mass per area (LMA).

#### Canopy ramification and wood density

We estimated the degree of canopy ramification as the number of terminal branch tips per stem cross-sectional area (BTSA; (Roddy *et al.*, 2019)) on plants growing in the common garden. Wood density of stems stripped of their bark was measured using Archimedes principle by measuring the mass of water displayed on a balance and subsequently measuring the dry mass of the stems.

#### Stem hydraulic conductance

Whole-shoot hydraulic conductance was measured using a low pressure flow meter (Kolb *et al.*, 1996), which enables measuring the entire shoot regardless of branch ramification and can be applied to morphologically diverse structures (Roddy *et al.*, 2016, 2019). Measurements were taken on healthy shoots from well-watered and mature plants. Hydraulic conductance was calculated as the slope of the regression of flow rate versus pressure. Because shoots differed in size and ramification, hydraulic conductance was normalized by leaf area of the shoot, which is taken as a metric of hydraulic efficiency (Roddy *et al.*, 2019).

#### MicroCT imaging

High-resolution, three-dimensional (3D) images of stem and leaf structure were obtained by performing hard X-ray microcomputed tomography (microCT) at the Advanced Light Source, Lawrence Berkeley National Laboratory (LBNL), Beamline 8.3.2 (Brodersen, 2013; Brodersen & Roddy, 2016). Full details of image capture are available in the Supplemental Information. Mature leaves and stems were sampled from plants growing in the common garden and imaged within 48 hours. Stems were allowed to air-dry prior to microCT imaging to ensure that vessels had emptied, and leaves were kept sealed in moist plastic bags until immediately before imaging.

To characterize trichome density, digital slices parallel to the fresh leaf surface were taken through the trichomes, allowing trichomes to be counted per unit projected leaf surface area. Stem xylem vessel diameter and area were measured using ImageJ on 2D cross sections of microCT image stacks obtained from dried stems.

### Spatial data collection

To determine the geographic extent and climatic envelope of *Encelia* species, we downloaded all occurrence data for *Encelia* species from the Global Biodiversity Information Facility (GBIF) on 10 June 2019 (GBIF, 2019). Using the R package CoordinateCleaner (Zizka *et al.*, 2019), we removed points falling in the ocean, zero coordinates, and equal coordinates and retained only those points from preserved specimens or human observations. By species, we then removed extreme spatial outliers using the `cc_outl` function, removed duplicate records, and thinned data by 1 kilometer using the R package spThin (Aiello-Lammens *et al.*, 2015).

Using these cleaned and thinned data, we extracted climatic and soil data using WorldClim 2.0 rasters at 30s resolution (Fick & Hijmans, 2017) and the Unified North American Soil Map at 0.25 degree resolution (Liu *et al.*, 2014). For climatic data, we focused on four bioclimatic variables that reflect extreme climatic conditions and are likely to be important in determining plant survival: maximum temperature of warmest month (bioclim 5), minimum temperature of coldest month (bioclim 6), precipitation of wettest month (bioclim 13), and precipitation of driest month (bioclim 14). We summarized the 18 soil variables describing soil composition and acidity using a principal component analysis (PCA) and retained the first two axes that explained 22% and 19% of the variation in total. All spatial analyses were conducted using the R packages raster and rgeos (Hijmans *et al.*, 2015; Bivand & Rundel, 2017).

### Comparative analyses

To determine net diversification rate, we used the crown age estimator across a range of extinction rates and our time-calibrated phylogeny (Magallón & Sanderson, 2001). We explored three scenarios for extinction (ε = 0.1, ε = 0.3, and ε = 0.9); the parameter ε reflects the balance between speciation and extinction rates.

To characterize the tempo and mode of trait evolution, we used species-level means and estimated phylogenetic signal (λ) for each trait (Pagel, 1999), calculated the rate of trait evolution in felsens (Ackerly, 2009), and calculated disparity through time using the average squared Euclidean distance (Harmon *et al.*, 2003). To measure correlations between traits and correlations between traits and environmental variables, we conducted phylogenetic generalized least squares (PGLS), using a Brownian motion correlation matrix. We used the R packages phytools, geiger, ape, nlme, and ggtree to conduct and visualize all comparative analyses (Paradis *et al.*, 2004; Harmon *et al.*, 2008; Pinheiro & J, 2009; Revell, 2012; Yu *et al.*, 2017).

To determine the biogeographic history of *Encelia*, we used BioGeoBEARS (Matzke, 2013) to estimate ancestral ranges. Occurrence data were mapped onto Ecological Regions of North America (Omernik & Griffith, 2014) to assign extant species to one or more of six biogeographic areas: Baja California Deserts, Mediterranean, Mojave Desert, Sonoran Desert, the cold deserts, or Peru (Fig. S1). We ran the Dispersal–Extinction–Cladogenesis (DEC) only, given limitations of more complex models (Ree & Sanmartín, 2018). We set the maximum possible range size to four.

We compared how species-pairs have diverged across environmental and morphological variables. For the morphological, soil, and climatic datasets, we first summarized the data using a scaled and centered principal component analysis (PCA). We then calculated species-level means across all PC axes; from these means, we calculated the Euclidean distance between all species-pairs in morphological and environmental space. For each pairwise comparison, we also determined geographic range overlap. To do this, we inferred geographic range for each species based on the alpha convex hull of species occurrence data and estimated overlap in convex hulls (Pateiro López & Rodríguez Casal, 2010).

## Results

### Genetic data analyses

Our final dataset resulted in an average of 870 Mb sequence across an average of 60K loci for 72 individuals (Table S1). We dropped five individuals that yielded <5% of homologous loci. Using these genetic data, we first determined likely lineage assignments among all sampled individuals, finding evidence for non-monophyly of *E. californica* and *E. virginensis* (Fig. S2). We accordingly revised lineage designations in these two nominal species to reflect the presence of putative new lineages (Table S1). The individual-level phylogeny based on these new lineage designations recovers the same topology as the lineage-level phylogeny (Figs. 1, 2) and the coalescent-based and concatenated phylogeny are largely concordant at the interspecific level (Fig. S2). The individual-level phylogeny exhibits high bootstrap support for the monophyly of all lineages but *E. palmeri* and *E. asperifolia* (Fig. 1). However, site-based and gene-level concordance (sCF and gCF) are low compared to bootstrap support (Fig. S3), which might be expected given the short internode distances in our phylogeny.

**Figure 2:**
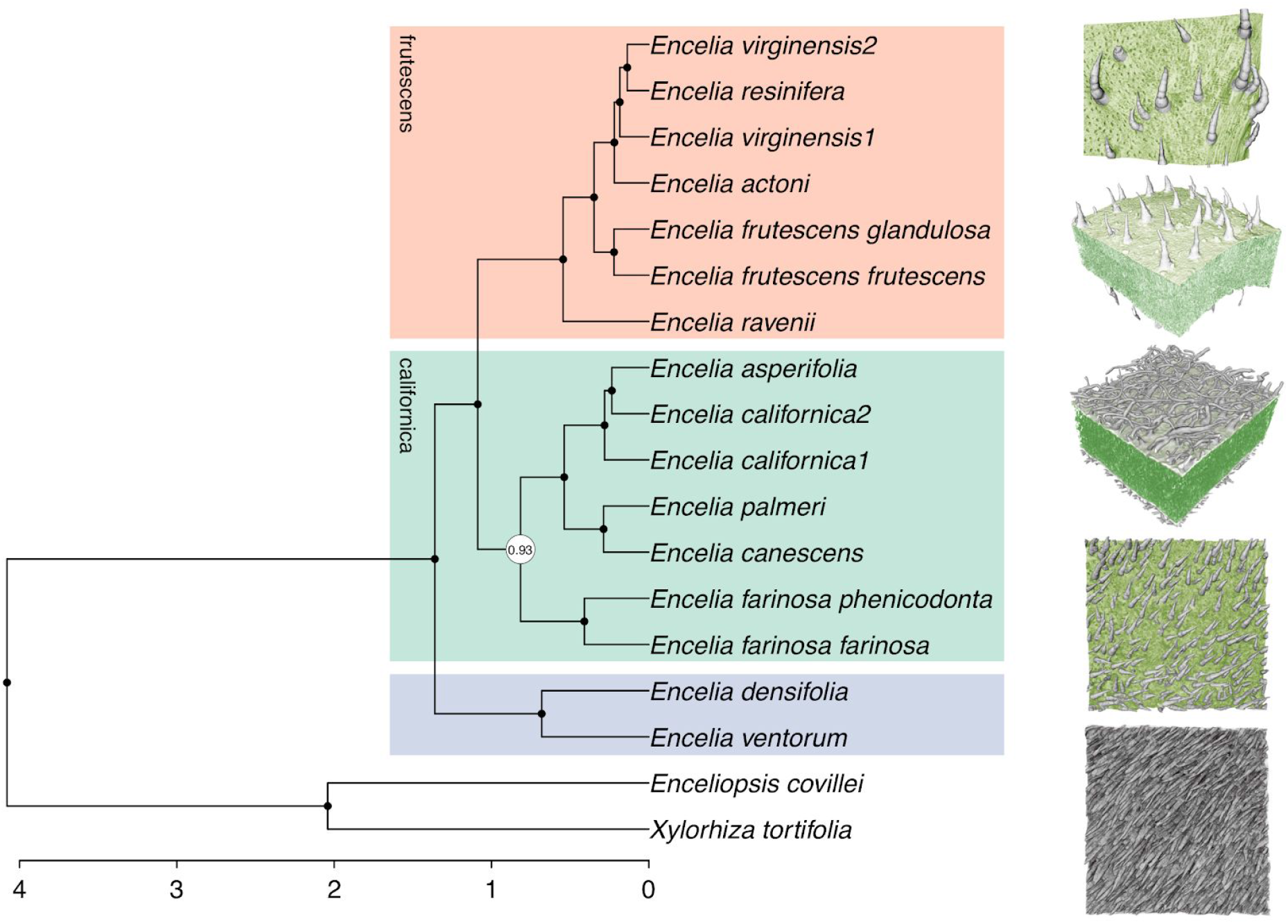
A lineage-level *Encelia* phylogeny inferred using the coalescent-based approach ASTRAL-III. MicroCT images show external leaf morphology of (top to bottom, with approximate widths of leading edges in parentheses): *E. frutescens frutescens* (950 μm), *E. asperifolia* (550 μm), *E. palmeri* (650 μm), *E. densifolia* (950 μm), and *Enceliopsis covillei* (475 μm). Images were false-colored to indicate green, photosynthetic tissue and how trichomes alter leaf color. Boxes demarcate major clades; time scale shown in millions of years. Nodes with less than 95% local posterior probability shown in white.

Both our lineage-level phylogenies were concordant (Fig. S4) and recovered three major clades (Fig. 2), two of which had been previously characterized as the *californica* and *frutescens* clades based on the species comprising the clades (Ehleringer & Clark, 1988; Fehlberg & Ranker, 2007). Additionally, we identified a third clade consisting of *E. densifolia* and *E. ventorum*. In contrast to previous phylogenetic studies for *Encelia*, statistical support for all nodes was uniformly high except for the placement of *E. farinosa*. The recent and rapid radiation in this group (Fig. 2) is also reflected in the low and relatively uniform levels of genetic divergence among lineages (average *F*_*ST*_ = 0.32, *D*_*xy*_ = 0.024, and *D*_*a*_ = 0.014, Fig. S5).

Phylogenetic networks inferred by SNAQ strongly supported one hybridization edge between *E. californica 2* and *E. asperifolia*, with admixture proportions of 0.49 and 0.51 between the two species (Fig. S6). Our D-statistic results identified multiple, strongly supported examples of introgression among lineages (Fig. S7; Table S3), including lineages that are known to hybridize in nature (e.g., *E. actoni* and *E. frutescens*; Fig. 3; Table S4).

**Figure 3:**
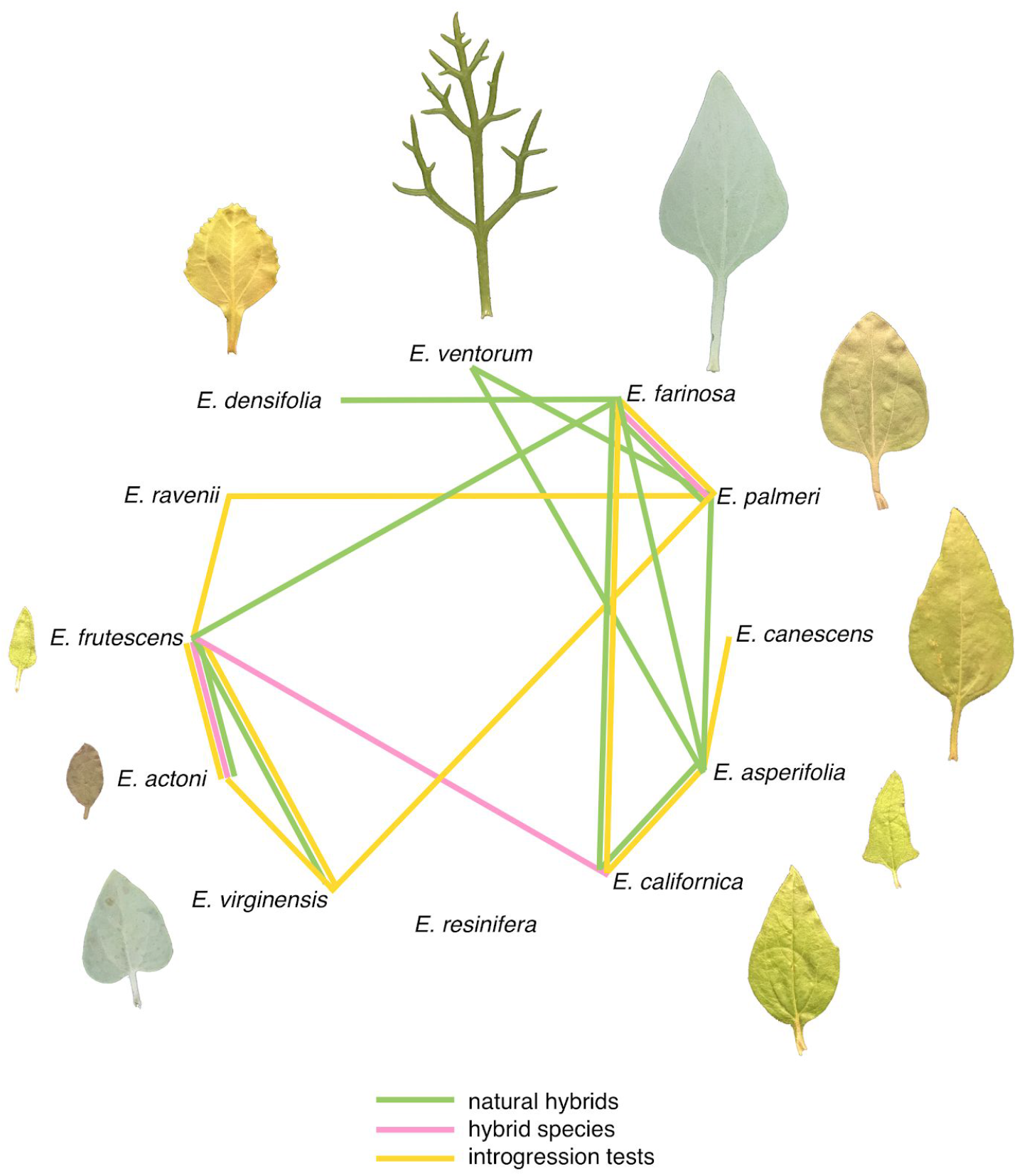
Hybridization and introgression in *Encelia* based on data from naturally-occurring hybrids, putative hybrid species, and tests of introgression (D-statistic & SNaQ analyses; Fig. S6, Fig. S7, Table S4). Species are arranged by clade identity and leaf images are relative to size. Hybridization and introgression are rampant across the clade, and many of the species-pairs show evidence for hybridization and introgression across multiple measures of introgression.

### Comparative analyses

Using the crown age estimator across a range of extinction rates and our time-calibrated phylogeny (Fig. 2), we found rates of diversification in *Encelia* vary from 1.57 (ε = 0.1), 1.52 (ε = 0.3), to 0.66 species per million years (Myr) (ε = 0.9) depending on the extinction scenario.

Reconstruction of trait evolution showed that closely-related species often have divergent trait values in nearly all phenotypes measured, indicative of widespread phenotypic divergence among *Encelia* species (Figs. 4, S8). This pattern of trait divergence is reflected in both low phylogenetic signal across all traits tested except leaf area (average lambda = 0.19, range = 0 to 1.16, Table 1) and rapid trait evolution (average felsens = 0.98, range = 0.02 to 6.23, Table 1). Rapid evolution in traits is mirrored by rapid transitions in environmental space in *Encelia*. Both within and among species and clades, a wide diversity of climatic space is occupied (Figs. S8, S9) and closely-related species often occupy very distinct climatic spaces (Figs. 4, S8). For example, individual species or clades can occupy temperatures from below freezing to >40°C or rainfall from close to 0 mm to 200 mm (Fig. S9).

**Table 1:** Estimates of phylogenetic signal (λ) and the associated significance and evolutionary rates (felsens; β) for each of the nine measured morphological and physiological traits.

| traits | $\lambda$ | p-val | $\beta$ | # of tips |
| --- | --- | --- | --- | --- |
| canopy ramification<br>(BTSA) | 0 | 1 | 1.17 | 13 |
| Trichome density (top) | 0 | 1 | 6.23 | 15 |
| Stem hydraulic<br>conductance | 0.4 | 0.83 | 0.57 | 8 |
| Leaf color | 0.15 | 0.84 | 0.03 | 11 |
| Leaf area | 1.16 | 0.01 | 0.49 | 11 |
| Leaf roundness | 0 | 1 | 0.04 | 11 |
| Leaf mass area (LMA) | 0 | 1 | 0.18 | 11 |
| Wood density | 0 | 1 | 0.02 | 5 |
| Vessel diameter | 0 | 1 | 0.09 | 11 |

**Figure 4:**
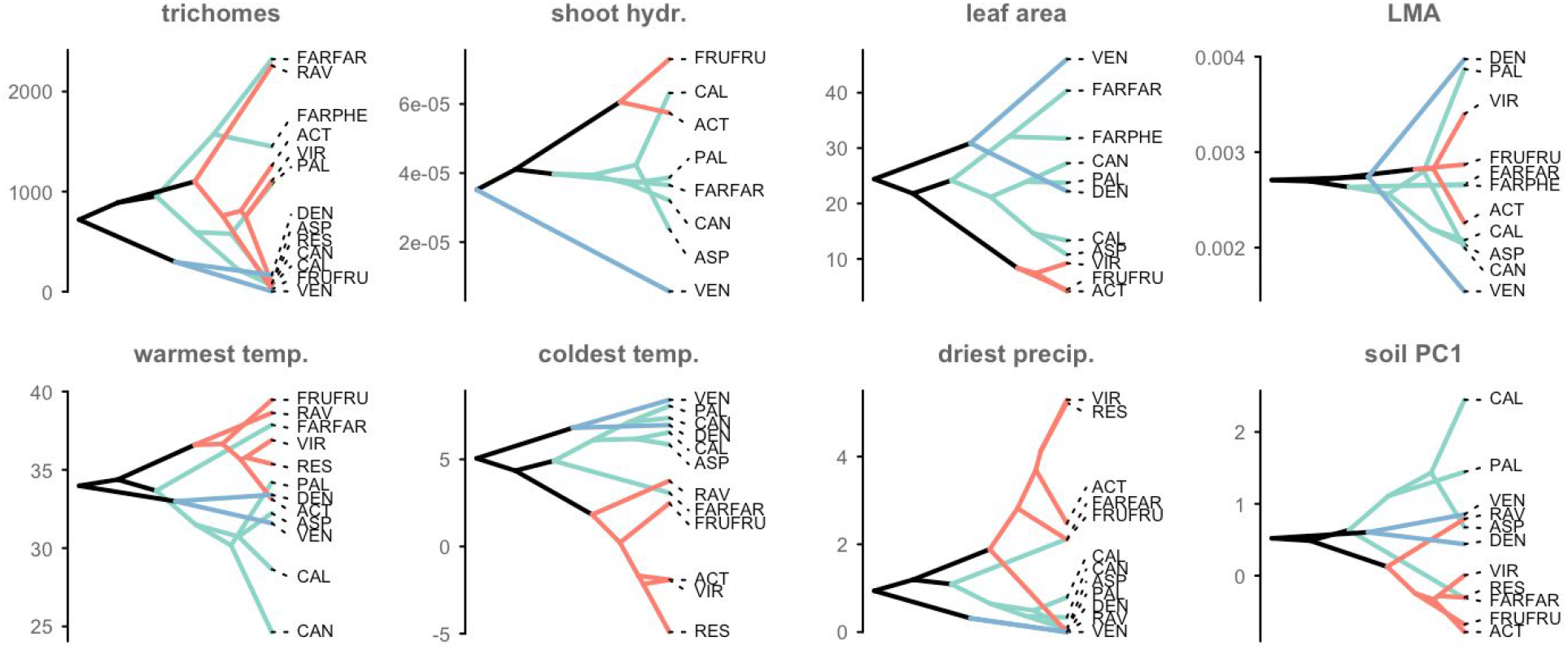
Phenotypic variation in *Encelia*, depicted as phenograms. The y-axis indicates phenotypic spread across (top) morphological and physiological traits and (bottom) environmental space. Abbreviations follow Table 1; all climatic variables are the extreme values across months. Branches are colored by clade identity as shown in Fig. 2, and all species names are abbreviated to the first three characters. Data for all traits and environmental measures shown in Fig. S8. Closely-related species in *Encelia* often exhibit dramatically different phenotypes.

Disparity-through-time (DTT) patterns fell within expectations under random Brownian evolution (Fig. S10). Across traits, correlations between traits were generally weak, and only a few trait combinations were significant (Fig. S11). Further, very few of the comparisons between traits and environmental variables were significant (Fig. S12), even though most correlations followed general physiological predictions (Table S5).

The ancestral range reconstruction under DEC in BioGeoBears returned fairly uncertain inference at deeper nodes. However, these results confirmed the origins of *Encelia* in some combination of the hot deserts (e.g., the Sonoran, Baja and Mojave Deserts; Fig. S1).

Following expectations, overlapping species-pairs showed greater similarity in climatic and soil envelopes than non-overlapping pairs (Fig. 5). However, many species that occupied the same environmental space -- including those that overlapped in geographic range -- differed as much in morphology as species pairs in completely different environmental spaces.

**Figure 5:**
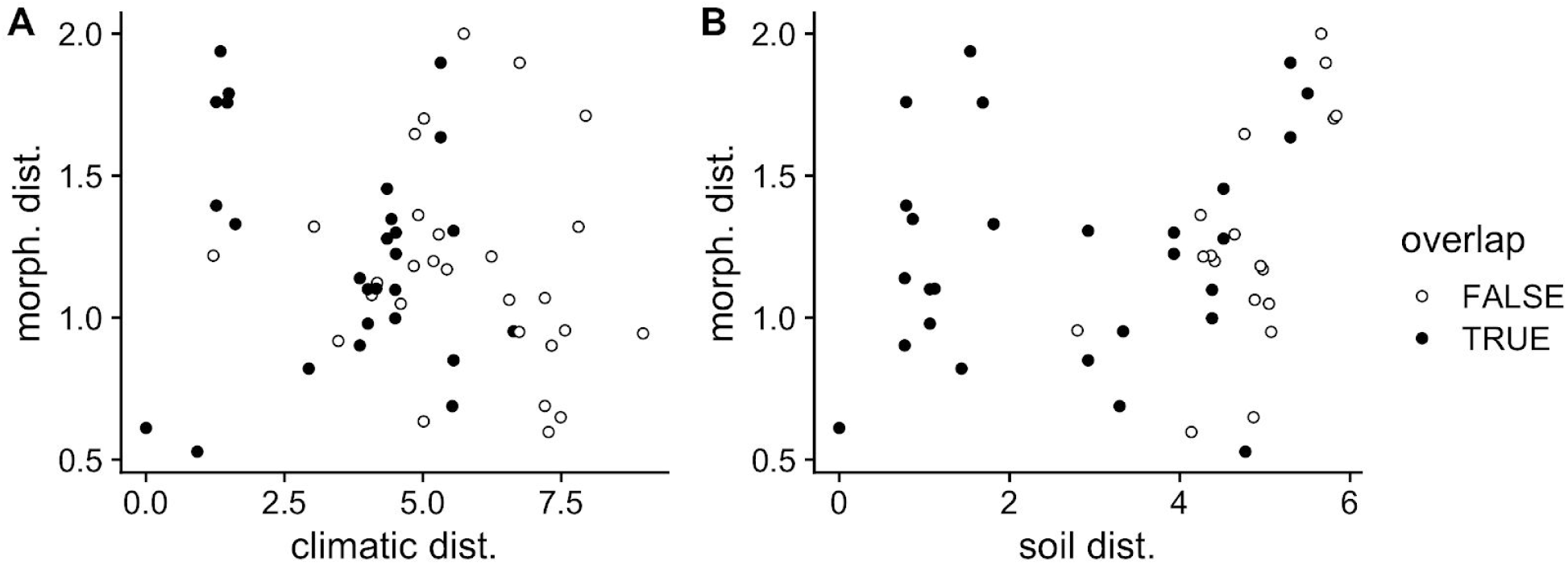
Pairwise species comparisons shown for (A) climatic distance between species versus morphological distance and (B) soil distance between species versus morphological distance. Comparisons are coded by whether or not their geographic ranges overlap. Many species-pairs occur in similar ecological environments (i.e., they exhibit low soil or climatic distance) but still exhibit high morphological distance.

## Discussion

*Encelia* is an enigmatic but charismatic group that has been a model system for ecophysiological studies of desert plants (Ehleringer *et al.*, 1981; Ehleringer, 1988; Ehleringer & Cook, 1990; Ehleringer & Sandquist, 2018). Previous phylogenetic studies based on both molecular and phenotypic data have failed to infer a well-resolved phylogeny of these plants because *Encelia* is a recent, rapid radiation and because of frequent hybridization (Fehlberg & Ranker, 2007). Aided by a phylogenomic dataset and extensive taxon sampling, we disentangled this radiation and resolved the evolutionary relationships among all sampled species (Fig. 2). In particular, we inferred the relationships between and within two previously identified clades: the *frutescens* clade, which includes the species found in the cold deserts of North America (*E. actoni, E. virginensis, E. resinifera, E. frutescens,* and *E. ravenii*), and the *californica* clade, which includes the majority of the diversity found in Baja California (*E. farinosa, E. canescens, E. palmeri, E. asperifolia, and E. californica*). Further, we found support for a new clade including *E. densifolia* and *E. ventorum*, both species restricted to Baja California. Together, these results suggest a pattern of phylogenetic eco-geographic structure whereby closely related species are largely restricted to the same or adjacent ecogeographic region (Figs. 1, S1).

### Rapid recent diversification and disparification in *Encelia*

*Encelia* has diversified recently and rapidly (Fig. 2), with 0.9 - 1.57 species produced per million years (Myr). In comparison, Hawaiian silverswords radiated at 0.56 species per Myr (Baldwin & Sanderson, 1998), Southern African ice plants at 0.77 - 1.75 species per Myr (Klak *et al.*, 2004), and New World *Lupinus* at 2.49 – 3.79 species per Myr (Hughes & Eastwood, 2006). Further, the two genera most closely related to *Encelia* (*Enceliopsis* and *Gerea*) are both relatively species-poor (four and two species, respectively), have older crown ages (Smith & Brown, 2018), and, accordingly, have much lower diversification rates. Thus, both within its local phylogenetic context and compared to other plant groups, *Encelia* has relatively high rates of diversification.

Concomitant with rapid diversification, *Encelia* shows rapid disparification. The absolute range of phenotypes seen within *Encelia* are broad even between closely related species, with leaf area varying 10-fold and trichome density varying >1000-fold (Figs. 4, S8). While this range is narrow relative to the full diversity seen in angiosperms (Wright *et al.*, 2004), it is striking given the young age of *Encelia*. Accordingly, *Encelia* has rates of trait evolution comparable to notable adaptive radiations. For example, leaf size evolves at an estimated 0.49 felsens compared to 0.46 in lobeliads and 2.08 in silverswords (Table 1, Ackerly, 2009). Unfortunately, we lack comparative data from other plant radiations for many of the traits we measured in *Encelia*. However, our estimated rates of evolution for other traits are high, exhibiting felsens >5 for trichome density, even without accounting for the multiple types of trichomes that occur among Encelia (Fig. 2; Ehleringer & Cook, 1987).

Although closely related species diverge extensively in phenotype, distantly related species often share similar phenotypes (e.g., *E. farinosa farinosa* and *E. ravenii* in trichome density, Fig. 4). This pattern of low phylogenetic signal suggests that accessibility to adaptive traits has enabled *Encelia* to diversify across a mosaic of environmental conditions and adaptive optima. Along with rapid trait evolution, *Encelia* species exhibit transitions along climate and soil gradients (Fig. 4, S8, S9), suggesting that *Encelia* species are capable of rapidly adapting to novel environmental conditions. Taken together, our findings suggest that high tait lability and rapid trait evolution are key syndromes underpinning the evolvability of the *Encelia* radiation.

### Drivers of diversification and disparification in *Encelia*

*Encelia* represents an excellent system for testing hypotheses for why deserts can generate exceptional diversity. First, the recent formation of arid habitats has provided new habitats in which desert-adapted species can diversify (Mooney & Zavaleta, 2016). Indeed, though reconstruction of *Encelia*’s ancestry ranges was inconclusive, it suggested *Encelia* most likely originated in the hot deserts, from which the *frutescens* clade spread into the cold deserts around ~0.5 myr (Fig. S1). Outside of the two species (*E. canescens* and *E. hispida*) that colonized Peru and the Galapagos Islands, the center of biodiversity of *Encelia* is in the deserts of North America. These deserts have changed through time and across space over the last 5 million years, with their initial formation and subsequent volcanic eruptions that likely eradicated much of the living flora (Conly *et al.*, 2005; Garrick *et al.*, 2009), sea incursions that divided the peninsula into isolated landmasses (Holt *et al.*, 2000; Riddle *et al.*, 2000), tectonic movement that led to the creation of the Baja peninsula (Dolby *et al.*, 2015), and glacial climate cycles that affected sea levels and habitat distributions (Van Devender & Spaulding, 1979; Thompson & Anderson, 2000). Although many phylogeographic studies place these landscape changes in the Pliocene, their timing is uncertain and certainly could have influenced the diversification of *Encelia* (Wilson & Pitts, 2010). These changes both allowed colonization of new habitats and divided existing populations, resulting in population isolation that would lead to increased diversification. Indeed, the population structure of multiple animal and plant species in the North American deserts (Riddle *et al.*, 2000; Crews & Hedin, 2006; Garrick *et al.*, 2009) reflects this history, most notably with splits across northern and southern Baja California and across the Sonoran and Baja Californian deserts. *Encelia* also presents evidence of this geographic pattern; of the five species that occur on the Baja peninsula, four of them (*E. densifolia*, *E. palmeri*, *E. ventorum* and *E. asperifolia*; Fig. 1) are restricted to the southern portion of the peninsula.

Second, although deserts are often characterized as homogenous swaths of arid land, deserts span large and often steep topological and environmental gradients, which can drive divergent selection and ecological speciation. Although all *Encelia* species are concentrated in North American deserts, they span multiple such gradients. *Encelia* species live in a diversity of climatic niches, experiencing hottest month temperatures ranging from 25 to 40°C, coldest month temperatures ranging from −5 to 8°C, and driest months ranging from 0 to 5 inches of rain (Figs. 4, S8). Although some of these absolute differences are small, they can represent large relative differences when resources are so limited. Notably, a few species are climatic outliers such as *E. californica*, which lives along the California coast, experiences high rainfall (Fig. S8, S9), and *E. actoni*, *E. resinifera*, and *E. virginensis* which all survive freezing temperatures (Fig. 4, S9). This climatic variance exemplifies the types of gradients that *Encelia* spans and that can drive diversification and disparification. Unlike species in many rapid radiations (Givnish, 1997), *Encelia* species are rarely sympatric and instead tend to share parapatric boundaries, defined by local environmental and edaphic transitions. Although contemporary geographic distributions do not necessarily reflect historical distributions, this pattern of species turnover across environmental gradients suggests that spatial environmental heterogeneity might have driven *Encelia* speciation.

Third, in arid habitats, multiple environmental gradients can interact and overlap at different spatial scales. Many *Encelia* species have parapatric geographic ranges (Fig. 1) and thus experience very similar climatic conditions (e.g., similar rainfall and solar insolation; Fig. 5). However, many of these species pairs are segregated along strong environmental gradients that occur over just a few meters, leading to marked phenotypic differences. Furthermore, the possible anatomical, physiological, and phenological adaptations to living in the stressful and resource-limited conditions of deserts are numerous, and combinations of these adaptive traits may all be equally fit, resulting in both high disparification and diversity (Stebbins, 1952; Roddy *et al.*, 2020). For example, *E. ventorum* occurs in sandy dunes that face the ocean, which encroach upon the inland deserts where *E. palmeri* is found (Kyhos *et al.*, 1981; DiVittorio *et al.*, 2020). *E. ventorum* can access the water table below the dunes but is also constantly exposed to osmotic stress from ocean spray. Thus, although these two species occur adjacent to each other and experience similar climates, they experience different levels of water availability and salt stress as well as different soil types, factors that together drive their phenotypic divergence in trichome density, leaf size, and shoot hydraulics (Fig. 2, 4). Similarly, *E. frutescens* and *E. farinosa* both occur in Death Valley, one of the hottest places in North America. *E. farinosa* has large leaves with abundant trichomes, while *E. frutescens* has small leaves with few trichomes (Fig. 4). These differences in leaf morphology influence their microhabitat occupations. *E. frutescens* occurs in wash habitats with higher water availability and uses transpirational cooling from its small leaves to maintain low leaf temperatures. In turn, *E. farinosa* occurs along dry slopes where its trichomes reflect solar radiation to maintain low leaf temperatures (Ehleringer, 1988). The same relationships between trichome density, leaf size, and water access also seem to influence microhabitat occupation by *E. palmeri* and *E. ventorum* (DiVittorio et al., 2020). These traits might have evolved repeatedly throughout *Encelia* in relation to fine-scale environmental heterogeneity.

These case studies exemplify how no single trait may be responsible for driving the radiation of *Encelia*. Rather, the high lability of multiple traits with compensatory physiological and fitness effects may have driven rapid trait evolution in *Encelia*, resulting in heightened diversification. As seen in other plant clades, trait lability could be a critical organismal syndrome to evolutionarily accessible phenotypes, which enables diversification within and across environments (Ogburn *et al.*, 2015). Writ large, trait lability would explain the lack of clear correlations between traits and broad-scale environmental conditions in *Encelia* (Fig. S12), weak correlations among traits (Fig. S11), and the presence of geographically-overlapping species that differ dramatically in morphology (Fig. 5). The diversity of traits associated with desert survival in *Encelia* exemplifies how aridity can be a catalyst of diversification and disparification.

### Hybridization and introgression and the *Encelia* radiation

In addition to biogeographic and environmental factors, hybridization and introgression have both contributed to the rapid diversification and disparification of *Encelia* and served as a source of genetic variation and for new species (Anderson & Stebbins, 1954; Stebbins, 1959; Marques *et al.*, 2019). Using two complementary approaches, we found numerous examples of introgression across *Encelia*, including across non-sister species and species in different major clades (Figs. S6, S7). In many cases, the instances of introgression are corroborated by field data of naturally occurring hybrids and by cases of suspected hybrid speciation (Fig. 3). Rampant hybridization has now been uncovered in multiple rapid radiations, such as the Hawaiian silversword alliance (Barrier *et al.*, 1999), African cichlids (Meier *et al.*, 2017), and *Heliconius* butterflies (Edelman *et al.*, 2019). To this list, we can now add *Encelia*.

Several aspects of *Encelia* biology and geography likely enabled this history of hybridization and introgression. *Encelia* species exhibit few of the barriers that restrict gene flow in other species: all species except *E. canescens* and likely *E. hispida* are obligate outcrossers, chromosome number is conserved across the genus, the species are pollinated by generalist pollinators, and, at the regional scale, they have only modest differences in flowering phenology (Ehleringer & Clark, 1988; Clark, 1998). Further, many *Encelia* species ranges are adjacent to each other (Fig. 1), across which dispersal is more permissible than barriers like mountains. Lastly, in the cold deserts, which are home to a number of species in the *frutescens* clade, midden data suggest that repeated glacial cycles led to repeated range retractions to relictual populations followed by range expansions (Spaulding & Graumlich, 1986; Thompson & Anderson, 2000). These recurrent bouts of secondary contact could drive introgression between species (Fehlberg & Ranker, 2009; Hewitt, 2011; Folk *et al.*, 2018), as outlined in the species-pump hypothesis (Papadopoulou & Knowles, 2015).

This history of hybridization and introgression might promote diversification by helping originate hybrid species. Previous studies of *Encelia* identified four putative cases of hybrid species (Table S4, Fig. 3): *E. actoni* x *E. frutescens* to result in both *E. virginensis* and *E. resinifera*, *E. californica* x *E. frutescens* to result in *E. asperifolia*, and *E. farinosa* x *E. palmeri* to result in *E. canescens* (Clark & Allan, 1997; Allan *et al.*, 1997; Clark, 1998). In each of these four cases, our D-statistic and SNAQ results confirm introgression edges either between the parental species or between the parental species and the putative hybrid species. Most notably, the admixture edge from *E. californica* to *E. asperifolia* was estimated at approximately 50%, as would be expected in hybrid speciation (Fig. S6). Confirming these cases of hybrid speciation will require more detailed analyses of the genome and spatial structure of these parental and putative hybrid species. Yet, these results provide compelling evidence that diversification in *Encelia* is partially driven by hybrid speciation.

Introgression can also help spur radiations by increasing the amount of genetic variation in a species (Suarez-Gonzalez *et al.*, 2018). Genetic variation can arise from a few sources: *de novo* mutations, standing genetic variation, and gene flow between populations or species. Typically, *de novo* mutations are thought unlikely to occur rapidly enough to drive rapid divergence (Barrett & Schluter, 2008). In contrast, both standing genetic variation and gene flow can provide an influx of variation to diverging populations (Hedrick, 2013; Suarez-Gonzalez *et al.*, 2018), allowing them to quickly adapt to new ecological conditions. In either case, distantly related species would exhibit similar phenotypes (Lee & Coop, 2019; Jamie & Meier, 2020), as occurs in *Encelia* (Fig. 4, S8). As yet, it is unclear if shared traits in *Encelia* arose from differential sorting of ancestral variation, introgression, or true convergent evolution. Reconstructing this history will require identifying the loci underpinning the trait of interest and characterizing their specific histories (Giska *et al.*, 2019).

Lastly, this history of introgression and hybridization in *Encelia* suggests that the genus might be a syngameon (Clark, 1998), a group of otherwise distinct species interconnected by gene exchange (Lotsy, 1925; Grant, 1971; Hipp *et al.*, 2019). In this scenario, these species can be fully independent evolutionary lineages that retain their cohesiveness and distinctiveness despite hybridization. Species participating in the syngameon can persist as a consequence of reinforcement, assortative mating, divergent selection, or selection against hybrids (Cannon & Petit, 2020). In particular, hybrid zone data suggest that extremely strong, divergent selection may be a predominant factor maintaining species boundaries in *Encelia* (DiVittorio *et al.*, 2020). Given that geographically overlapping species often are quite phenotypically divergent (Fig. 5), divergent selection might be helping to maintain species cohesiveness throughout the genus.

## Conclusion

Our integrative study within a single desert lineage provides new insights into the processes of plant evolution in one of the harshest terrestrial environments. The evolutionary history of *Encelia* provides an example of a radiation encompassing rapid and recent species formation, high phenotypic disparity, and strong ecological divergence - thus meeting many of the key requirements of an adaptive radiation (Givnish, 1997). Eco-phenotypic differentiation that results in functional and fitness tradeoffs are apparent across this radiation. Rather than homogenizing genetic lineages, interspecific gene flow increases genetic diversity within species and may facilitate adaptation. We suggest that the combined effects of high genetic diversity with high trait lability have enabled access to multiple adaptive peaks, leading to species diversification and disparification across steep environmental gradients both at broad and fine spatial scales. Much remains to be learned about the mechanisms underpinning this radiation. The patchy environmental heterogeneity characteristic of the deserts presents an exciting opportunity to model explicitly the influence of a complex fitness landscape with multiple optima on the genomic background of a radiating lineage (Martin & Richards, 2019). The extent to which interspecific gene flow has enabled adaptation in this scenario is a critical area for continued study.

## Acknowledgements

We acknowledge funding from UC Mexus to CD & SS, from a NSF Postdoctoral Fellowship in Biology to SS (DEB-1519732), a CSUDH Norris Faculty Grant to SS, and by a postdoctoral fellowship from the Yale Institute for Biospheric Studies to ABR. We thank the UC Riverside AgOps team for their efforts in maintaining the common garden and Danny Eduardo Carvajal López and Darren Sandquist for providing seeds for *E. canescens*. Comments from I. Holmes improved a previous version of this manuscript. MicroCT imaging was performed at the Lawrence Berkeley National Laboratory Advanced Light Source Beamline 8.3.2 microtomography facility, with help from D. Parkinson and A. MacDowell. The Advanced Light Source is supported by the Director, Office of Science, Office of Basic Energy Services, of the U.S. Department of Energy under contract no. DE-AC01-05CH11231.

## Author Contributions

SS, ABR, FZ designed the research. SS, ABR, CD, CLH, CRB, ASA, and FZ conducted the research. SS, ABR, and FZ analyzed and interpreted data. SS, ABR, and FZ wrote the manuscript.

## Data Availability

Raw sequence data are available at SRA <accession numberTBD>. Alignments and variant call sets used in population genetic and phylogenetic inference are available on DataDryad <DOI TBD>. Trait measurements are available on DataDryad <DOI TBD>. All scripts used to analyze and visualize data are available on GitHub <link TBD>.

## Details on Supplemental Information

*Expanded Methods:* Expanded details on how genetic, trait, and spatial data were collected and analyzed in this study.

*Figure S1:* Ancestral range reconstruction in *Encelia* based on the DEC model implemented in BioGeoBEARS.

*Figure S2:* Individual-level phylogenies for *Encelia* inferred using a concatenated maximum-likelihood approach and a coalescent-based approach.

*Figure S3:* Nodal support for the individual-level phylogeny for Encelia as measured by bootstrap values and site and gene concordance factors.

*Figure S4:* Lineage-level phylogenies for *Encelia* inferred using a concatenated maximum-likelihood approach and a coalescent-based approach.

*Figure S5:* Genetic divergence between *Encelia* species, as measured by FST, dxy, and da.

*Figure S6:* Reconstruction of *Encelia*’s evolutionary history as a network using SNaQ.

*Figure S7:* Introgression across *Encelia* as measured by the D-statistic.

*Figure S8:* Phenotypic variation in *Encelia*, depicted as phenograms.

*Figure S9:* The environmental space occupied by *Encelia* species across climatic variables.

*Figure S10:* Disparity-through-time plots for phenotypic variation in morphological traits and environmental space.

*Figure S11:* Pairwise correlations among all measured traits in *Encelia*.

*Figure S12:* Correlations between trait and environmental variables in *Encelia*.

*Table S1:* Detailed information on the 72 individuals included in this study, including their locality and species designations.

*Table S2:* The nine morphological and physiological traits measured in this study.

*Table S3:* Results of significant D-statistic tests for introgression.

*Table S4:* Instances of hybridization and introgression in the *Encelia* genus from both this study and previously-collected data.

*Table S5*: Expected correlations between traits and different environmental variables and the results recovered in this study.

