## Supplementary Information for "Diversification, disparification, and hybridization in the desert shrubs *Encelia*"

### Supplementary Methods

#### Sampling: Common Garden

Collected seeds were planted in a common garden for use in all genetic and phenotypic analyses. The common garden was first planted in 2011 and then expanded in 2016. By using a common garden to collect both genetic and phenotypic data, we ensured that our phenotypic data were matched to a genotype and that phenotypic differences among individuals were not due to environmental variation. Seeds were germinated in greenhouse trays using an equal mix of pumice, sand, and vermiculite. *Encelia ravenii* and *E. resinifera* were not included in the garden because their seeds failed to germinate. When seedlings were 5 cm tall or had four true leaves, they were transplanted into the common garden at the University of California Agricultural Operations Station in Riverside, CA. Seedlings were randomized across rows and then irrigated weekly. Plants were phenotyped from 2015 to 2016, and in October 2016, adult leaves were collected in silica gel for use in genetic analysis. For phenotypic measurements of *E. ravenii* and *E. resinifera*, we used field-collected adult leaves.

#### Genetic data collection

Because ddRAD data have not yet been collected for *Encelia*, we first conducted a series of *in silico* experiments to determine the best restriction enzymes and target size to use. The closest available reference genome to *Encelia* is *Helianthus annuus*, another species of the family Asteraceae estimated to be 7.5 million years diverged from *Encelia* (Smith & Brown, 2018). Many of the species in Asteraceae, including *Encelia*, have extensive repetitive content (Badouin *et al.*, 2017). To avoid sequencing this repeat content, we focused on testing restriction enzymes that are sensitive to the DNA methylation common in repeat content (Pegadaraju *et al.*, 2013). In total, we tested 14 different restriction enzymes in all possible pairwise combinations. Ultimately, to give the desired yield of 20,000 loci, we used *PstI* and *MspI*, size-selecting between 250 - 700 bp.

To collect ddRAD data, we first extracted DNA from silica-dried adult leaves using a QIAGEN DNeasy Plant Kit (cat. no.: 69104) following manufacturer's instructions. We determined DNA quality by running extracts on a 2% agarose gel and DNA concentration by using a NanoDrop DNA spectrometer. The commercial provider RTL Genomics (Lubbock, TX, USA) then prepared doubly-barcoded ddRAD libraries (Peterson *et al.*, 2012). Equimolar amounts of samples were combined to generate pools of 100 individuals; each pool was sequenced across one lane of 100 PE sequencing on the Illumina HiSeq 4000 Sequencing Platform.

Per individual, we first cleaned all reads, trimming adaptors and low quality sequence using `Trimmomatic v36` (Bolger *et al.*, 2014). We then merged any overlapping reads using `PEAR v0.9.8` (Zhang *et al.*, 2014). Due to extensive adaptor contamination, many of our paired reads became orphaned through the cleaning process. We then assembled cleaned and merged reads using `Velvet v1.2.10` (Zerbino & Birney, 2008) across multiple k-mer values. Although `Velvet` is not often used in ddRAD assembly, other commonly used tools like `Rainbow` or `STACKS` are not designed to assemble mixed sets of single-end and paired-end reads. We then retained all assembled loci with coverage ≥2×, resulting in an average of 140K loci per individual.

**Phylogenetic Inference**

Although all sampled individuals were identified to nominal species, species boundaries have not been well-tested in *Encelia*. Thus, our first set of analyses were designed to determine lineage identity per individual. We used `VSEARCH v2.9.1` (Rognes *et al.*, 2016) to identify homologous loci using a 70% identity cutoff. After aligning homologous loci using `mafft v7.9.34` (Katoh *et al.*, 2009), we then concatenated homologous loci across individuals, removing any individual sequenced for <5% of loci or any loci sequenced for <30% of individuals. We used `RAxML v8.2.11` to generate a phylogeny from the 11.6K locus, 1.5 Mb concatenated alignment (Stamatakis, 2014). By comparing clade identity to nominal species identity, we determined likely lineage identities for each individual (Table S1).

We then inferred individual-level phylogenies using both concatenated and coalescent-based approaches. First, we used the lineage-level pseudo-reference genomes and the variant call set to determine loci sequences per individual. We removed any sequences with >30% of missing sites and any loci sampled for <60% of individuals. We then used `RAxML` to infer a phylogeny and 100 bootstraps from the 31K locus, 3.9 Mb concatenated alignment. For a coalescent-based approach, we used `SVDquartets` (Chifman & Kubatko, 2014). We randomly sampled one SNP per locus, retaining SNPs with <20% missing and ≥2 minor allele count. We used the resulting set of 2.8K SNPs to infer the phylogeny and to conduct 100 bootstrap analyses. Bootstrap estimates of nodal support are often inflated, particularly for phylogenomic datasets (Cummings *et al.*, 2003). Accordingly, we calculated gene concordance factors (gCF) and site concordance factors (sCF) using `IQ-TREE v1.6.4` (Minh *et al.*, 2020), both of which better measure conflict across loci and sites.

Then, we used two different coalescent-based approaches - one based on gene trees and the other based on SNPs - to infer a lineage-level phylogeny. We used `ASTRAL-III` (Zhang *et al.*, 2018), which, when given an individual-to-lineage mapping and a set of gene trees inferred across individuals, can generate a lineage-level phylogeny. Here, we filtered loci to include only loci inferred for >60% of individuals, inferred gene trees using `RAxML`, collapsed all randomly-bifurcated nodes using 'dimulti' in the R package `ape` (Paradis *et al.*, 2004), and then inferred the species tree from these gene trees with `ASTRAL-III`. Second, using the same SNP dataset used for the individual-level `SVDquartets` analysis, we additionally inferred a lineage-level phylogeny. Neither `ASTRAL-III` nor `SVDquartets` provides terminal branch lengths. Accordingly, to infer an ultrametric phylogeny for use in comparative analyses, we generated a lineage-level concatenated alignment by randomly selecting one sequence per locus per lineage. We then used `RAxML` to estimate branch lengths, constraining the topology to each of the lineage-level phylogenies. To infer an ultrametric tree for this group, we used a calibration from a comprehensive angiosperm phylogeny that estimated the crown age of *Encelia* as 1.36 million years (Myr) (Magallón *et al.*, 2015; Smith & Brown, 2018). This aligns with previous divergence dating based on population genomic data that inferred the root of *Encelia* as 1.05 Myr (Singhal, unpublished). We used this root age to infer an ultrametric tree using the 'chronos' function in the R package `'ape'` with lambda of 0.01.

### MicroCT imaging

High-resolution, three-dimensional (3D) images of stem and leaf structure were obtained by performing hard X-ray microcomputed tomography (microCT) at the Advanced Light Source, Lawrence Berkeley National Laboratory (LBNL), Beamline 8.3.2 (Brodersen, 2013; Brodersen & Roddy, 2016). These images were used to visualize fine-scale morphology and anatomy of these leaves and to calculate trichome density and stem xylem vessel diameter and area. Leaf samples were collected from plants growing in cultivation at the Agricultural Operations Station, University of California, Riverside in June 2015 and September 2016 and transported to LBNL, keeping the leaves hydrated until imaging. Samples were placed on a rotating stage in the 24 keV synchrotron X-ray beam and continuously rotated from 0 to 180 degrees. As the sample rotated continuously through the beam, 1025 two-dimensional projections were recorded using a 10x objective, yielding a final pixel resolution of 1.25 µm. Each scan was completed in ~15 min. These raw tomographic projections were reconstructed using TomoPy, a Python-based framework for reconstructing tomographic data (Gürsoy *et al.*, 2014). Images were reconstructed using gridrec (Dowd *et al.*, 1999) and phase retrieval (Davis *et al.*, 1995) reconstruction methods. Image stacks were rotated, aligned, and traits measured in FIJI (Schindelin *et al.*, 2012). Three-dimensional stack visualization was performed in Avizo v. 9.2.0 (ThermoFisher Scientific).

### Trait data collection & analysis

#### *Leaf area, shape, and color*

To analyze leaf area, shape, and color, we collected three to five adult leaves per individual growing in the common garden. Using a digital camera, we photographed these leaves with a black-and-white reference scale bar. Leaves were then oven-dried at 70 C for at least 72 hours and weighed for dry mass using a Sartorius Practum (resolution of 0.0001 mg, Sartorius AG, Goettingen, Germany). We analyzed leaf images using ImageJ (Rasband & Others, 1997), first calibrating the white balance and size of each image using the scale bar as a reference (Mascalchi, 2017). For each leaf, we set the threshold across the leaf, excluding the petiole, and then collected data on area, perimeter, fit ellipse, and shape using the Analyze Particles function. To measure color, we used the white-balanced leaf images in Adobe Photoshop and measured the arithmetic mean of all pixels of the largest circumscribed rectangle possible within the center of the leaf. We calculated leaf mass per area (LMA) using measurements of leaf area and dry mass. We conducted a principal component analysis of all leaf size and shape measurements after centering and scaling them. The first two axes summarize 42% and 24% of the variation respectively; PC1 largely loads on size-related measurements and PC2 largely reflects the roundness of the leaf.

*Canopy ramification*

We estimated the degree of canopy ramification as the number of terminal branch tips per stem cross-sectional area (BTSA; Roddy *et al.*, 2019). For each plant in the common garden, we took two perpendicular measurements of the diameter of the stem base using manual calipers and averaged these two measurements to estimate the area of the stem base. We then counted all branch tips descendant from this base, regardless of their size.

*Trichome density*

Trichomes were characterized from 3D microCT images of fresh leaves. Subregions of each leaf surface were cropped and rotated to align the leaf surface with the plane of view. Digital slices parallel to the leaf surface were taken through the trichomes, allowing trichomes to be counted per unit projected leaf surface area.

*Vessel dimensions*

Stem xylem vessel dimensions were measured from 2D cross sections of microCT image stacks obtained from dried stems. Images were thresholded and binarized to distinguish vessel lumens from plant tissue, and the area of each of these lumens measured using the Analyze Particles function in ImageJ.

*Wood density*

Wood density was measured using Archimedes' principle. A small segment of stem was excised, its bark removed, and its volume measured as the mass of water displaced when it was submerged in a beaker of water sitting on a balance. The segment was then oven-dried at 70 C for at least one week, after which its dry mass was measured.

*Stem hydraulic conductance*

Whole-shoot hydraulic conductance was measured using a low pressure flow meter (Kolb *et al.*, 1996), which enables measuring the entire shoot regardless of branch ramification and can be applied to morphologically diverse structures (Roddy *et al.*, 2016, 2019). Mature plants growing under well-watered conditions at the Agricultural Operations Station, University of California, Riverside, were sampled in May 2013. Healthy shoots were selected from each plant.

In the late evening, entire shoots were cut at their base, and immediately recut underwater while the shoot was enclosed in black plastic bags and transported to the lab. Shoots were allowed to rehydrate for at least two hours before hydraulic measurements, and measurements were completed within 24 hours of sampling. Branching shoots were defoliated underwater, the leaves scanned for leaf area, and the cut shoot base was recut under clean water with a fresh razor blade. While keeping the cut stem surface covered in water, the shoot base was quickly enclosed in a compression fitting that was connected to a hard-sided tube, which was filled with and plumbed back to a bottle of 0.01 M KCl that sat on a microbalance with resolution to 0.0001 g (Sartorius CPA225D, Sartorius AG, Goettingen, Germany). The shoot was enclosed in an acrylic vacuum chamber. The vacuum chamber was connected to a pressure gauge and a vacuum pump, which created a series of partial vacuums (60, 50, 40, 30, 20, 15 kPa below ambient) on the stem that pulled KCl from the balance and into the stem. Every fifteen seconds the mass of KCl on the balance was logged by a computer interfaced with the balance (WinWedge, TAL Technologies Inc., Philadelphia, PA, USA). The partial vacuum was maintained for at least three minutes at each pressure and until the coefficient of variation of the previous ten measurements was below 0.05. The average of these last ten measurements was taken as the flow rate at the given pressure. Hydraulic conductance was calculated as the slope of the regression of flow rate versus pressure, and no more than one of the six measurements for each shoot was removed to improve linearity. Because shoots differed in size and ramification, hydraulic conductance was normalized by leaf area of the shoot, which is taken as a metric of hydraulic efficiency (Roddy *et al.*, 2019).

### Supplementary Tables

**Table S1:** Information on the 72 individuals included in this study, including their species identification, the locality from which they were collected, and whether they are in our common garden. For those nominal species that contain multiple putative lineages, we indicate lineage identity in parentheses. This table also outlines the efficacy of our genetic data collection, including trimmed sequencing read yield in megabases (Mb), number of assembled homologous loci, and number of high-quality, high coverage variant and invariant sites.

| sample | species | locality | In common garden | read yield (Mb) | loci count | sites (Mb) | SRA |
|---|---|---|---|---|---|---|---|
| ACT-9M-1 | *Encelia actoni* | 9 Mile Canyon Road, CA, USA | * | 2055.59 | 68861 | 12.62 | TBD |
| ACT-9M-2 | *E. actoni* | 9 Mile Canyon Road, CA, USA | * | 223.83 | 37805 | 5.29 | TBD |
| ACT-AB-1 | *E. actoni* | Anza Borrego, CA, USA | * | 176.34 | 33263 | 4.78 | TBD |
| ACT-AB-2 | *E. actoni* | Anza Borrego, CA, USA | * | 102.6 | 22625 | 3.17 | TBD |
| ASP-CD-1 | *E. asperifolia* | Central Desert, BC, Mexico | * | 72.55 | 19214 | 2.27 | TBD |
| ASP-CD-2 | *E. asperifolia* | Central Desert, BC, Mexico | * | 308.75 | 37977 | 5.46 | TBD |
| ASP0129 | *E. asperifolia* | 10 km south of El Rosarito, BC, Mexico | | 87.05 | 20879 | 2.45 | TBD |
| CAL-CR-1 | *E. californica (1)* | north of El Rosario, BC, Mexico | * | 79.68 | 17522 | 2.21 | TBD |
| CAL-CR-2 | *E. californica (1)* | north of El Rosario, BC, Mexico | * | 64.54 | 13257 | 1.72 | TBD |
| CAL-SC-1 | *E. californica (1)* | San Clemente, CA, USA | * | 177.14 | 23995 | 3.38 | TBD |
| CAL-SC-2 | *E. californica (1)* | San Clemente, CA, USA | * | 214.8 | 27656 | 4.25 | TBD |
| CAL-ELR-1 | *E. californica (2)* | pass south of El Rosario, BC, Mexico | * | 72.35 | 13977 | 1.86 | TBD |
| CAL-ELR-2 | *E. californica (2)* | pass south of El Rosario, BC, Mexico | * | 78.58 | 17616 | 2.24 | TBD |
| CAN-COP-5 | *E. canescens* | Atacama Desert, Chile | | 384.14 | 40606 | 6.47 | TBD |
| CAN-DOM-1 | *E. canescens* | Atacama Desert, Chile | | 483.94 | 13355 | 1.07 | TBD |
| CAN-HUA-4 | *E. canescens* | Atacama Desert, Chile | | 494.51 | 42540 | 7.26 | TBD |
| CAN-HUA-6 | *E. canescens* | Atacama Desert, Chile | | 325.6 | 27905 | 4.76 | TBD |

| | | | | | | | |
|---|---|---|---|---|---|---|---|
| CAN-ING-1 | *E. canescens* | Atacama Desert, Chile | | 122.41 | 14309 | 2.31 | TBD |
| CAN-OBI-7 | *E. canescens* | Atacama Desert, Chile | | 330.76 | 32032 | 5.26 | TBD |
| CAN-OBI-9 | *E. canescens* | Atacama Desert, Chile | | 85.97 | 18302 | 2.26 | TBD |
| CAN-PERU-2 | *E. canescens* | Arequipa 2, Peru | * | 4091.65 | 49865 | 11.81 | TBD |
| CAN-PERU-3 | *E. canescens* | Mollebaya, Peru | * | 1162.26 | 67133 | 8.96 | TBD |
| CAN-VAL-9 | *E. canescens* | Atacama Desert, Chile | | 769.06 | 62575 | 8.03 | TBD |
| DEN-12-A | *E. densifolia* | Sierra Santa Clara, BCS, Mexico | * | 513.66 | 55393 | 6.67 | TBD |
| DEN-13-A | *E. densifolia* | Sierra Santa Clara, BCS, Mexico | * | 323.22 | 52031 | 6.17 | TBD |
| DEN-16 | *E. densifolia* | Sierra Santa Clara, BCS, Mexico | * | 966.03 | 77650 | 10.23 | TBD |
| DEN-17 | *E. densifolia* | Sierra Santa Clara, BCS, Mexico | * | 279.31 | 50346 | 6.03 | TBD |
| DEN-18-A | *E. densifolia* | Sierra Santa Clara, BCS, Mexico | * | 365.55 | 52963 | 6.49 | TBD |
| DEN-21 | *E. densifolia* | Sierra Santa Clara, BCS, Mexico | * | 42.47 | 16433 | 1.44 | TBD |
| FAR-FAR-AB-1 | *E. farinosa farinosa* | Anza Borrego, CA, USA | * | 1186.43 | 90436 | 12.04 | TBD |
| FAR-FAR-AB-2 | *E. farinosa farinosa* | Anza Borrego, CA, USA | * | 1905.66 | 96822 | 13.27 | TBD |
| FAR-FAR-DVHZ-1 | *E. farinosa farinosa* | Death Valley, CA, USA | * | 815.27 | 79334 | 10.01 | TBD |
| FAR-FAR-DVHZ-2 | *E. farinosa farinosa* | Death Valley, CA, USA | * | 1381.38 | 87741 | 11.83 | TBD |
| FAR-FAR-DVSC-1 | *E. farinosa farinosa* | Surprise Canyon, Death Valley, CA, USA | * | 514.03 | 70868 | 8.93 | TBD |
| FAR-FAR-HH-1 | *E. farinosa farinosa* | Hidden Hills, Mojave National Preserve, CA, USA | * | 419.46 | 60795 | 7.14 | TBD |
| FAR-FAR-HH-2 | *E. farinosa farinosa* | Hidden Hills, Mojave National Preserve, CA, USA | * | 1324.88 | 88721 | 11.87 | TBD |
| FAR-FAR-RV-1 | *E. farinosa farinosa* | Riverside, CA, USA | * | 356.7 | 65021 | 7.91 | TBD |
| FAR-FAR-RV-2 | *E. farinosa farinosa* | Riverside, CA, USA | * | 1258.34 | 87994 | 11.57 | TBD |
| FAR-PHE-52 | *E. farinosa phenicodonta* | Central Desert, BC, Mexico | * | 675.51 | 77547 | 9.94 | TBD |
| FAR-PHE-60-2 | *E. farinosa phenicodonta* | Sea of Cortez, BC, Mexico | * | 892.79 | 77349 | 9.88 | TBD |
| FAR-PHE-71-1 | *E. farinosa phenicodonta* | north of San Felipe, BC, Mexico | * | 865.67 | 84450 | 11.11 | TBD |
| FAR-PHE-73-1 | *E. farinosa phenicodonta* | north of San Felipe, BC, Mexico | * | 696.14 | 71950 | 9.2 | TBD |

| | | | | | | | |
|---|---|---|---|---|---|---|---|
| FAR-SON | *E. farinosa phenicodonta* | Sonora, Mexico | * | 548.82 | 70368 | 8.6 | TBD |
| FRU-FRU-DV-1 | *E. frutescens frutescens* | Death Valley, CA, USA | * | 416.43 | 61365 | 7.45 | TBD |
| FRU-FRU-DV-2 | *E. frutescens frutescens* | Death Valley, CA, USA | * | 456.33 | 59658 | 7.13 | TBD |
| FRU-FRU-DVHZ-1 | *E. frutescens frutescens* | Death Valley, CA, USA | * | 1019.08 | 76237 | 9.68 | TBD |
| FRU-FRU-DVHZ-2 | *E. frutescens frutescens* | Death Valley, CA, USA | * | 683.58 | 70228 | 8.81 | TBD |
| FRU-FRU-MNP-1 | *E. frutescens frutescens* | Mojave National Preserve, CA, USA | * | 814.49 | 77813 | 9.78 | TBD |
| FRU-FRU-MNP-2 | *E. frutescens frutescens* | Mojave National Preserve, CA, USA | * | 1115.06 | 85680 | 11.03 | TBD |
| FRUGLA1 | *E. frutescens glandulosa* | 25 km north of Condensadora, BC, Mexico | | 1570.74 | 94144 | 12.62 | TBD |
| FRUGLA2 | *E. frutescens glandulosa* | 25 km north of Condensadora, BC, Mexico | | 462.31 | 69256 | 8.24 | TBD |
| PAL-BA-1 | *E. palmeri* | Bahia Asuncion, BCS, Mexico | * | 59.69 | 20398 | 1.81 | TBD |
| PAL-BA-2 | *E. palmeri* | Bahia Asuncion, BCS, Mexico | * | 1305.15 | 90331 | 12.08 | TBD |
| PAL-JESUS-01 | *E. palmeri* | Villa Jesus Maria, BC, Mexico | | 903.89 | 79699 | 10.43 | TBD |
| PAL648 | *E. palmeri* | El Marasal, BC, Mexico | | 1143.91 | 87751 | 11.71 | TBD |
| RAV-1 | *E. ravenii* | San Felipe Desert, BC, Mexico | | 1763.21 | 85833 | 11.45 | TBD |
| RAV-10 | *E. ravenii* | San Felipe Desert, BC, Mexico | | 1971.25 | 92493 | 12.62 | TBD |
| RAV-11 | *E. ravenii* | San Felipe Desert, BC, Mexico | | 1224.63 | 72385 | 9.89 | TBD |
| RAV-13 | *E. ravenii* | San Felipe Desert, BC, Mexico | | 1577.5 | 87206 | 11.73 | TBD |
| RAV-2 | *E. ravenii* | San Felipe Desert, BC, Mexico | | 1231.7 | 84033 | 11.05 | TBD |
| RES-1 | *E. resinifera* | Zion National Park, Washington Co., UT, USA | | 919.69 | 91989 | 12.08 | TBD |
| RES-2 | *E. resinifera* | Zion National Park, Washington Co., UT, USA | | 1322.5 | 98559 | 13.05 | TBD |
| VEN-BA-1 | *E. ventorum* | Bahia Asuncion, BCS, Mexico | * | 1452.92 | 89500 | 11.71 | TBD |
| VEN-BA-2 | *E. ventorum* | Bahia Asuncion, BCS, Mexico | * | 2350.32 | 97483 | 13.31 | TBD |

| | | | | | | | |
|---|---|---|---|---|---|---|---|
| VIR-GMDRC | *E. virginensis (1)* | Granite Mountains Desert Research Center, CA, USA | * | 1575.73 | 96090 | 12.92 | TBD |
| VIR-MNP-1 | *E. virginensis (1)* | Foshay Pass, Mojave National Preserve, CA, USA | * | 871.66 | 84453 | 11.29 | TBD |
| VIR-MNP-2 | *E. virginensis (1)* | Foshay Pass, Mojave National Preserve, CA, USA | * | 1792.58 | 98577 | 13.66 | TBD |
| VIR-UT-1 | *E. virginensis (2)* | Highway 91, Washington Co., UT, USA | | 1565.36 | 108361 | 14.61 | TBD |
| VIR-UT-2 | *E. virginensis (2)* | Highway 91, Washington Co., UT, USA | | 637.33 | 71703 | 9.24 | TBD |
| ENC-1 | *Enceliopsis covillei* | Surprise Canyon, Death Valley, CA, USA | * | 1066.09 | 48713 | 5.99 | TBD |
| ENC-2 | *Enceliopsis covillei* | Surprise Canyon, Death Valley, CA, USA | * | 1378.21 | 49010 | 6.34 | TBD |
| XYL | *Xylorhiza tortifolia* | Mojave National Preserve, CA, USA | * | 2756.57 | 10268 | 0.93 | TBD |

**Table S2:** The nine morphological and physiological traits measured in this study.

| trait | description | # of lineages measured |
|---|---|---|
| canopy ramification | measured as the number of branch tips per stem cross-sectional area (BTSA) | 15 |
| trichome density | top and bottom trichome density was measured on microCT leaf images; top and bottom trichome density were highly correlated ($r$ = 0.99, p-val < 5e-14) | 15 |
| stem hydraulic conductance | slope of flow rate versus pressure for stems, normalized by leaf area of shoot | 8 |
| leaf color | leaf color was calculated on color-balanced leaves as the arithmetic mean of pixels | 12 |
| leaf roundness | leaf roundness is highly correlated with leaf PC2 ($r$ = -0.98) which captures 24% of the variation | 12 |
| leaf area | leaf area is highly correlated with leaf PC1 ($r$ = -0.97, p-val < 2e-16) which captures 42.5% of the variation | 12 |
| leaf mass area (LMA) | leaf dry mass divided by leaf area | 12 |
| wood density | woody stem (stripped of bark) dry mass divided by its volume | 5 |
| vessel diameter | stem xylem vessel diameter | 11 |

**Table S3:** Results of significant D-statistic tests. Shown are D-statistic analyses that significantly deviate from 0, as measured by their Z-score. The two species inferred to have exchanged genes, as determined by the sign of the D-statistic, are listed in the first two columns.

| species 1 | species 2 | D-statistic | Z-score | p-val |
|---|---|---|---|---|
| *E. actoni* | *E. virginensis 1* | 0.19 | 34.66 | 3.12E-263 |
| *E. actoni* | *E. frutescens frutescens* | 0.06 | 8.52 | 1.62E-17 |
| *E. virginensis 1* | *E. frutescens frutescens* | 0.04 | 6.5 | 7.96E-11 |
| *E. californica 1* | *E. californica 2* | 0.1 | 5.13 | 2.86E-07 |
| *E. palmeri* | *E. farinosa farinosa* | 0.11 | 7.07 | 1.55E-12 |
| *E. palmeri* | *E. farinosa phenicodonta* | 0.03 | 6.17 | 6.79E-10 |
| *E. palmeri* | *E. ravenii* | 0.03 | 4.62 | 3.83E-06 |
| *E. ravenii* | *E. frutescens frutescens* | 0.03 | 5.44 | 5.40E-08 |
| *E. frutescens frutescens* | *E. virginensis 2* | 0.04 | 6.97 | 3.06E-12 |
| *E. frutescens glandulosa* | *E. virginensis 2* | 0.07 | 10.73 | 7.73E-27 |

**Table S4:** Instances of hybridization and introgression in the *Encelia* genus. Shown are the involved species, what type of data was used to infer the hybridization or introgression event, and notes, including the reference for the data. Data types "dstat" and "SNaQ" come from the present study.

| species 1 | species 2 | data type | notes |
|---|---|---|---|
| *E. actoni* | *E. frutescens* | natural hybrids | no explicit data, Clark 1998 |
| *E. actoni* | *E. frutescens frutescens* | dstat | conservative test |
| *E. actoni* | *E. frutescens frutescens* | hybrid species | result in *E. virginensis*; genetic analysis from Allan et al 1997 |
| *E. actoni* | *E. frutescens glandulosa* | hybrid species | result in *E. resinifera*; no formal test but included in Allan et al. 1997 |
| *E. actoni* | *E. virginensis* 1 | dstat | conservative test |
| *E. asperifolia* | *E. californica* | natural hybrids | field data from DiVittorio et al., unpublished |
| *E. asperifolia* | *E. californica* 2 | SNaQ | approximately 50% admixture |
| *E. asperifolia* | *E. canescens* | dstat | conservative test |
| *E. asperifolia* | *E. farinosa* | natural hybrids | no explicit data, Clark 1998 |
| *E. asperifolia* | *E. palmeri* | natural hybrids | field data from DiVittorio et al., unpublished |
| *E. asperifolia* | *E. ventorum* | natural hybrids | Kyhos et al. 1981; DiVittorio et al. 2020; Clark 1998 |
| *E. californica* | *E. farinosa* | natural hybrids | http://nathistoc.bio.uci.edu/plants/Asteraceae/Encelia%20hybrids/Encelia%20hybrids.htm |
| *E. californica* | *E. frutescens* | hybrid species | result in *E. asperifolia*; no formal test but included in Allan et al. 1997 |
| *E. californica* 1 | *E. californica* 2 | dstat | conservative test |
| *E. californica* 2 | *E. farinosa* | dstat | conservative test |

| | | | |
|---|---|---|---|
| | *phenicodonta* | | |
| *E. densifolia* | *E. farinosa* | natural hybrids | field data from DiVittorio et al., unpublished |
| *E. farinosa* | *E. frutescens* | natural hybrids | field data from DiVittorio et al., unpublished; Clark 1998 |
| *E. farinosa* | *E. palmeri* | hybrid species | result in *E. canescens*; no formal test but included in Allan et al. 1997 |
| *E. farinosa* | *E. palmeri* | natural hybrids | no explicit data, Clark 1998 |
| *E. farinosa farinosa* | *E. palmeri* | dstat | conservative test |
| *E. farinosa phenicodonta* | *E. palmeri* | dstat | conservative test |
| *E. frutescens* | *E. virginensis* | natural hybrids | no explicit data, Clark 1998 |
| *E. frutescens frutescens* | *E. ravenii* | dstat | conservative test |
| *E. frutescens frutescens* | *E. virginensis* 1 | dstat | conservative test |
| *E. frutescens frutescens* | *E. virginensis* 2 | dstat | conservative test |
| *E. frutescens glandulosa* | *E. virginensis* 2 | dstat | conservative test |
| *E. palmeri* | *E. ravenii* | dstat | conservative test |
| *E. palmeri* | *E. ventorum* | natural hybrids | Kyhos et al. 1981; DiVittorio et al. 2020; Clark 1998 |
| *E. palmeri* | *E. virginensis* 2 | dstat | conservative test |

**Table S5:** Expected correlations between traits and different environmental variables and the correlation recovered in this study, and whether or not it is significant (sig.) or not (n.s.). Abbreviations follow Table 1.

| trait | environmental variable | predicted correlation | found correlation |
| --- | --- | --- | --- |
| trichomes | precipitation | - | +, n.s. |
| trichomes | temperature | + | +, n.s. |
| leaf area | precipitation | + | +, n.s. |
| leaf area | temperature | - | -, n.s. |
| LMA | precipitation | + | +, n.s. |
| LMA | temperature | + | +, sig. |

### Supplementary Figures

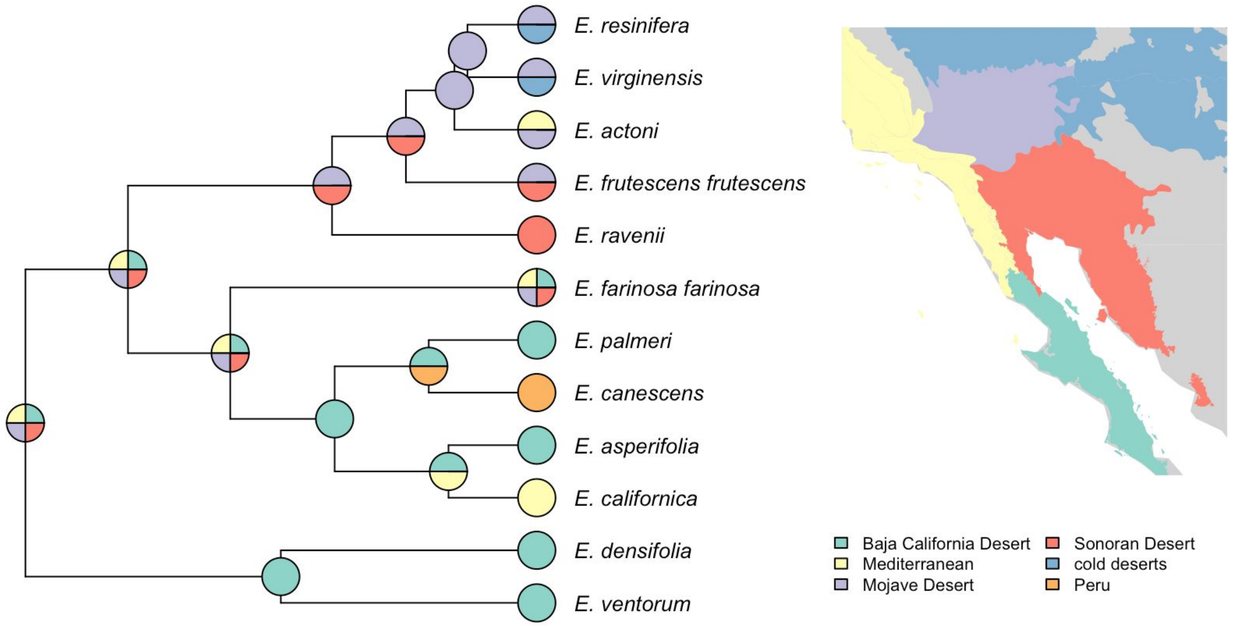


**Figure S1:** Ancestral range reconstruction in *Encelia* based on the DEC model implemented in `BioGeoBEARS`. Left shows phylogeny; pie charts at tips show the biogeographic ranges to which species were assigned and pie charts at nodes represent the ancestral range reconstruction with the highest probability at that node. The map shows the biogeographic regions.

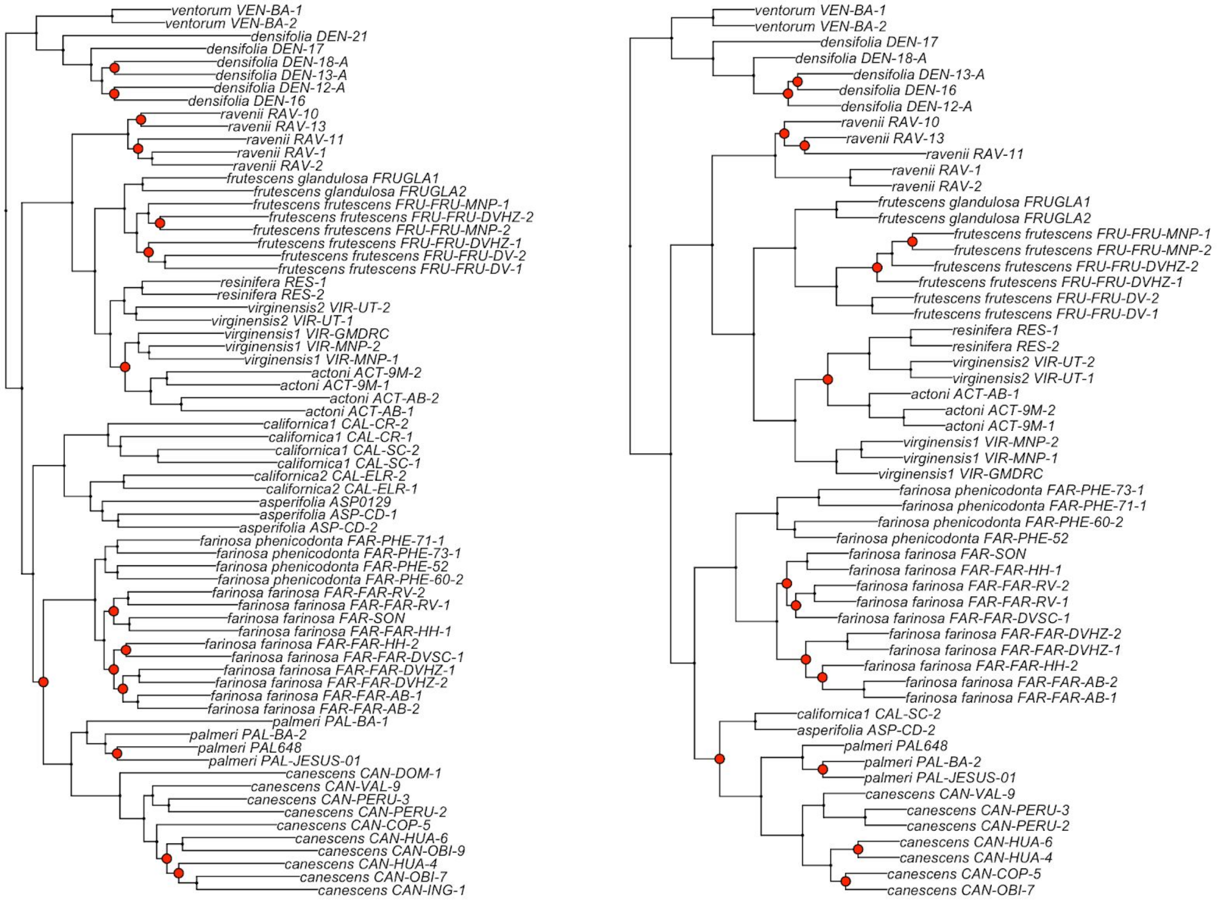


**Figure S2:** Individual-level phylogenies for *Encelia* inferred using (L) a concatenated maximum-likelihood approach implemented in RAxML and (R) a single nucleotide polymorphism (SNP) coalescent-based approach implemented in SVDquartets. The SVDquartets phylogeny (n = 56) contains a subset of the tips in the RAxML phylogeny (n = 69) because individuals with high levels of missing SNP data were removed. Phylogenies rooted by outgroups (not shown). Nodes labeled in red differ between the two topologies. Most of the nodes that differ are among individuals within a species.

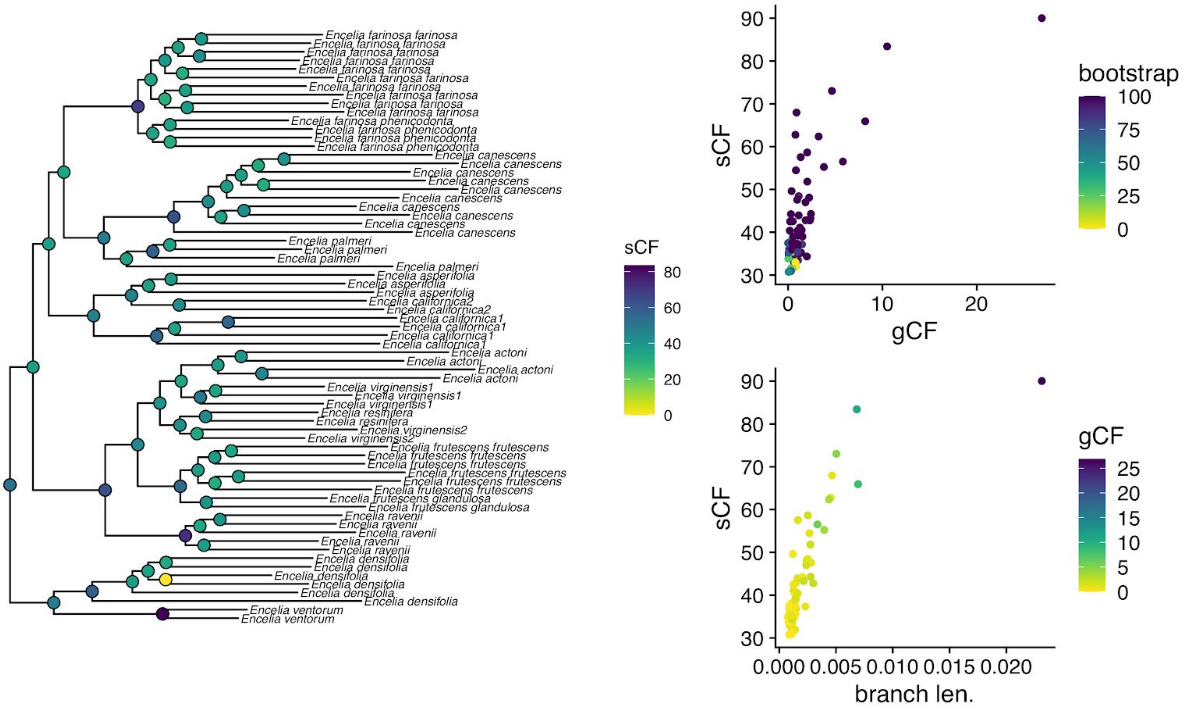


**Figure S3:** Individual-level phylogenies for *Encelia* inferred using a concatenated maximum-likelihood approach implemented in `RAxML`. Phylogeny rooted by outgroups (not shown). Nodal values indicate site concordance factors (sCF). Plots show (top) gene concordance factors (gCF) relative to sCF and (bottom) sCF relative to branch length. sCF and gCF values are relatively low throughout the topology. Low sCF and gCF values are expected in recent rapid radiations due to extensive incomplete lineage sorting.

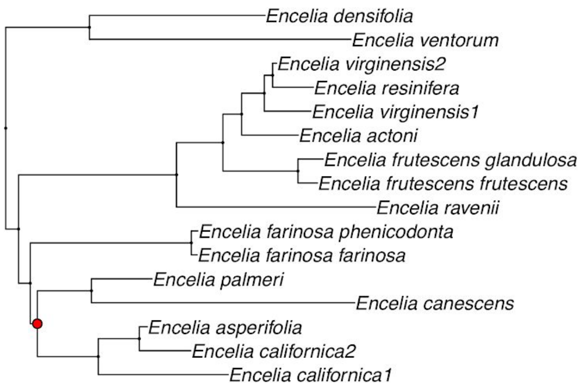


**Figure S4:** Lineage-level phylogenies inferred using coalescent-based approaches (L) using gene trees implemented in `ASTRAL` and (R) single nucleotide polymorphisms implemented in `SVDquartets`. Nodes that differ between the two topologies shown in red. The sole node that differs between the two topologies has low local posterior probability (0.93; Fig. 2).

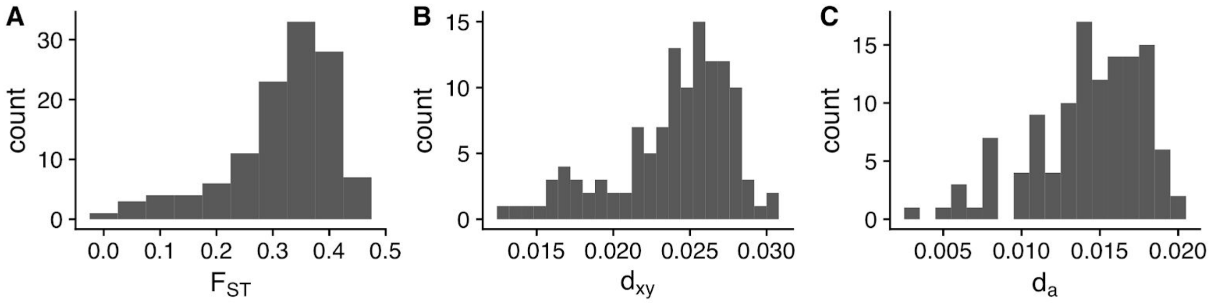


**Figure S5**: Genetic divergence between *Encelia* species, as measured by (A) $F_{ST}$, (B) $d_{xy}$, and (C) $d_a$. As expected under a recent, rapid radiation, genetic divergences across species-pairs are both low and similar.

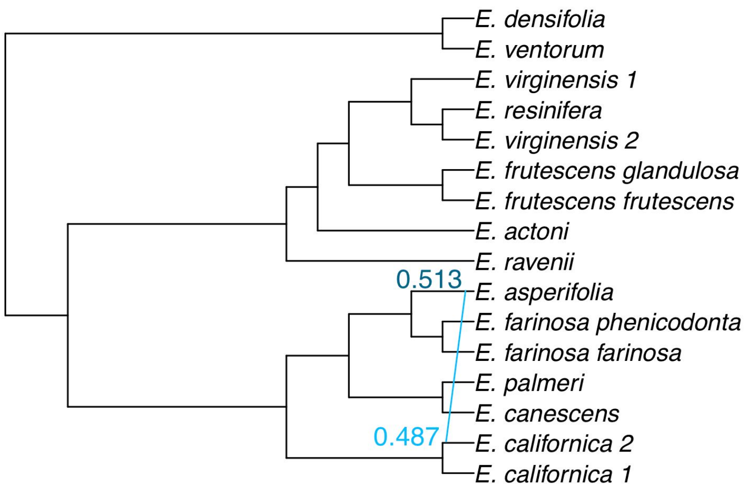


**Figure S6:** Reconstruction of *Encelia*'s evolutionary history as a network using `SNaQ`. The best fit model inferred one reticulation event; *E. asperifolia* is a hybrid with ~50% ancestry of *E. californica 2*. *Encelia asperifolia* has been hypothesized to be a hybrid species resulting from hybridization between *E. californica* and *E. frutescens glandulosa* (Table S4), although this hypothesis has not been formally tested previously.

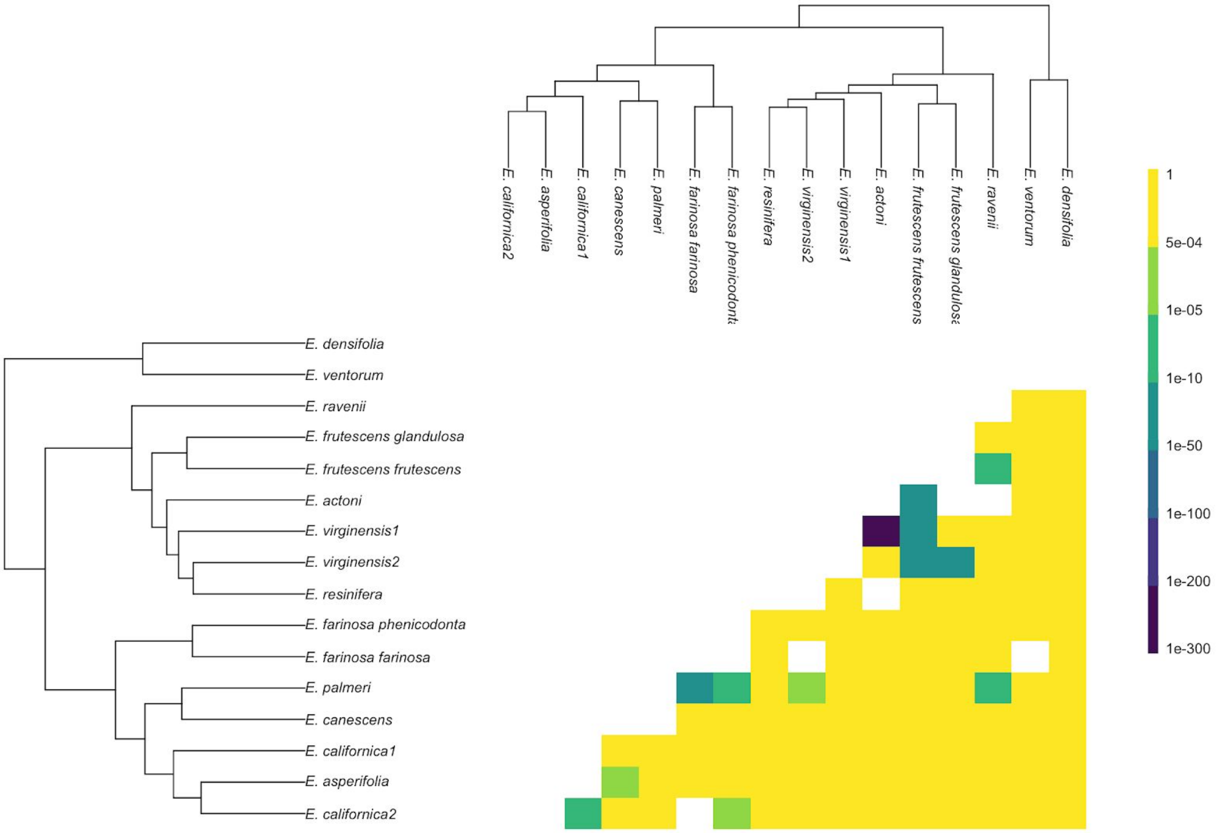


**Figure S7:** Introgression across *Encelia* as measured by the D-statistic. Plotted are the p-values of measured D-statistics (Table S3). The D-statistic was calculated across all possible triads in *Encelia*; for a given species pair, shown is the greatest p-value. Insignificant tests are shown in yellow; all significant tests are shown in shades of green & blue. Some species pairs are shown in white. For these species pairs, we either could not evaluate the D-statistic because they are sister species or because, in all comparisons involving these two species, introgression occurred with the third species. Even using this conservative approach, several species-pairs show strong evidence for introgression, including some species known to hybridize in nature (Fig. 3).

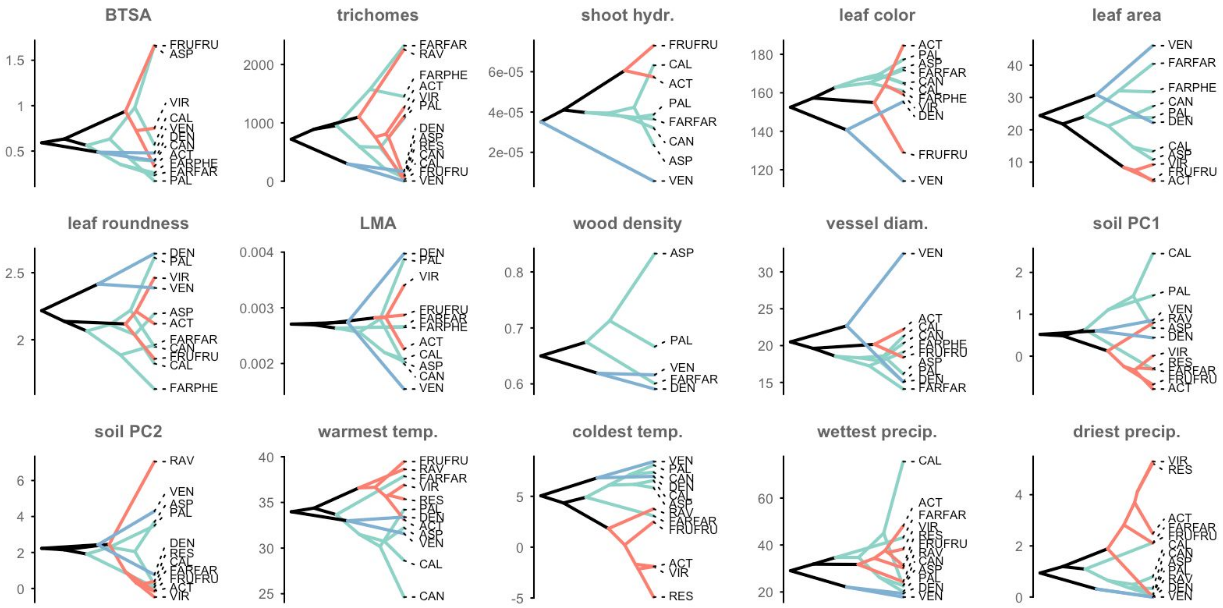


**Figure S8:** Phenotypic variation in *Encelia*, depicted as phenograms. The y-axis indicates phenotypic spread across (top two rows) morphological and physiological traits and (bottom row) environmental variation. Trait abbreviations follow Table 1. Branches are colored by clade identity as shown in Fig. 2, and all species names are abbreviated to the first three characters. Closely-related species in *Encelia* often exhibit dramatically different phenotypes.

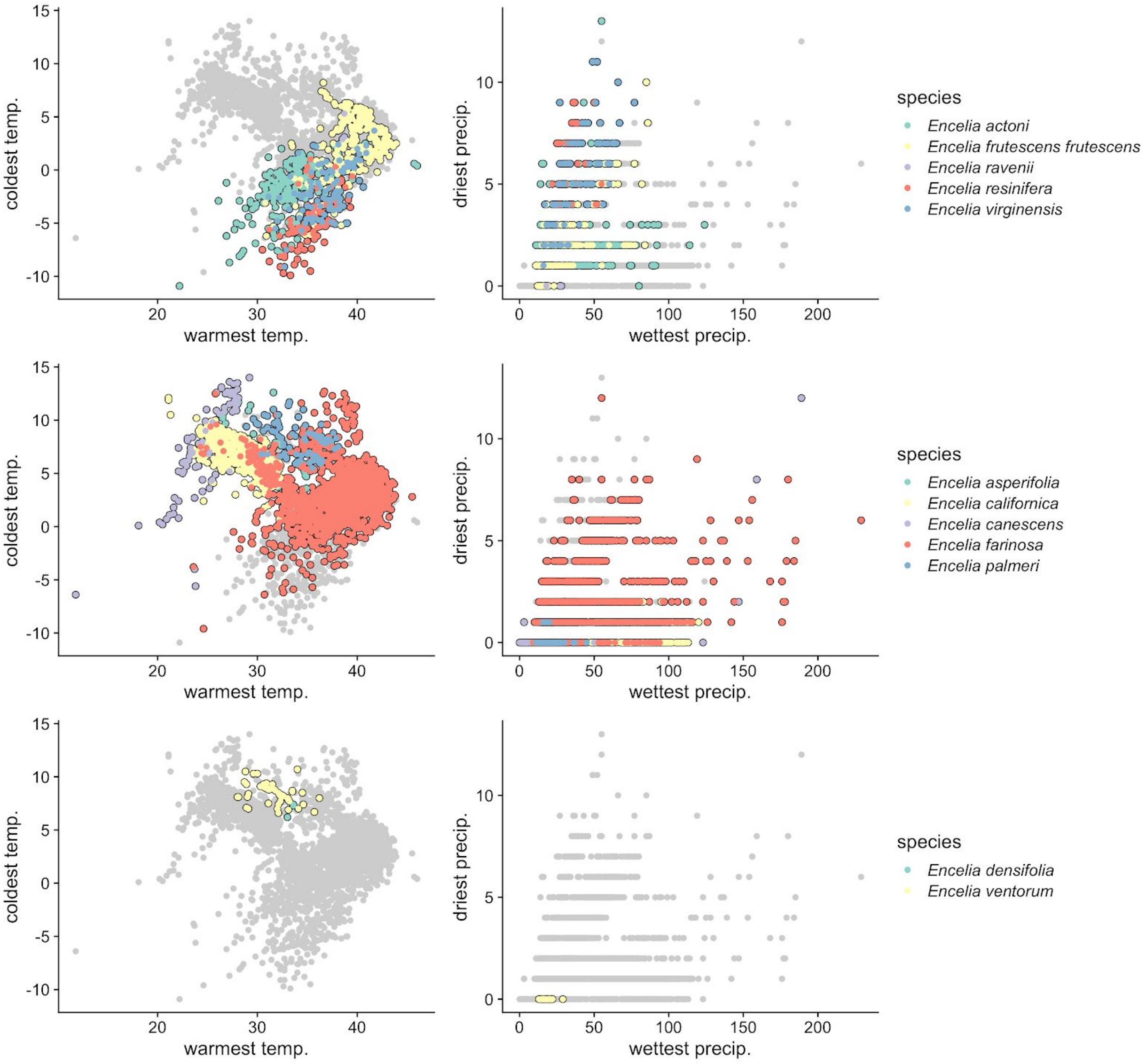


**Figure S9:** The environmental space occupied by *Encelia* species, as summarized by four bioclimatic variables: maximum temperature of the warmest month (BIO5), minimum temperature of the coldest month (BIO6), precipitation of wettest month (BIO13), and precipitation of driest month (BIO14). Species are broken up by the major clades in which they occur (Fig. 2). The species in *Encelia* occupy a diversity of climatic environments, although many of them overlap in climatic space.

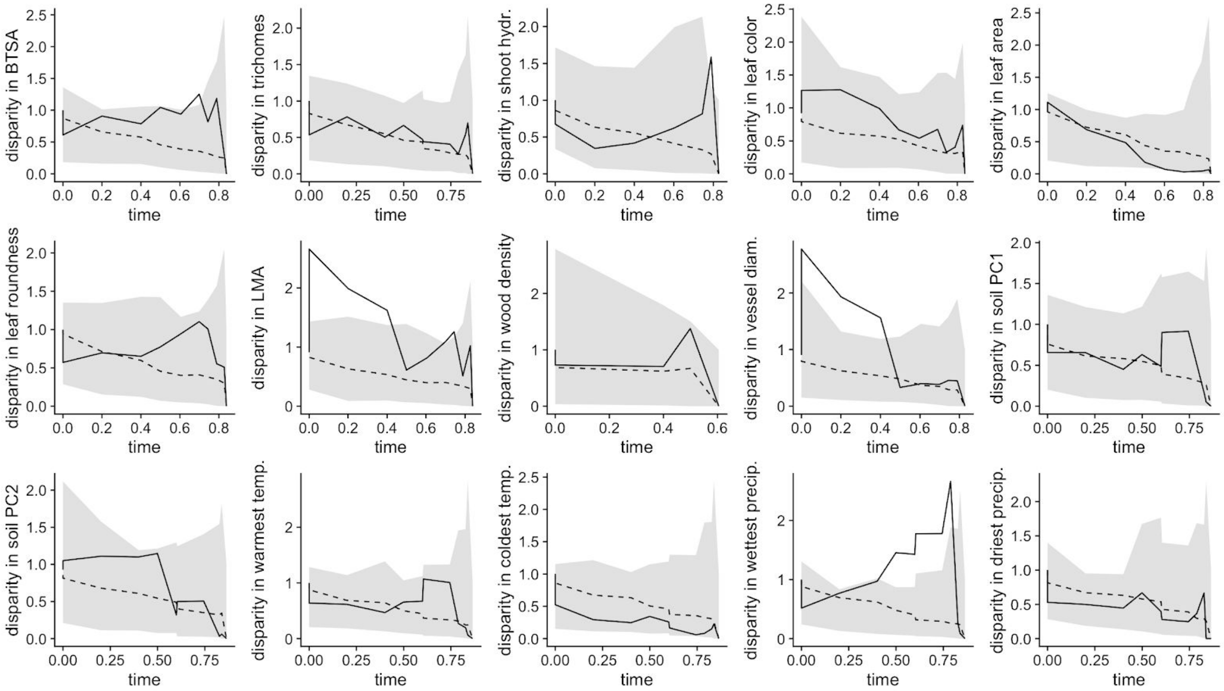


**Figure S10:** Disparity-through-time plots for phenotypic variation in morphological traits and environmental space. Solid black lines show disparity as measured in *Encelia*. The gray area indicates the 95% confidence interval across 100 simulations and the dotted black lines indicate the mean across Brownian motion simulations. Unlike many adaptive radiations, *Encelia* does not show evidence of early disparification.

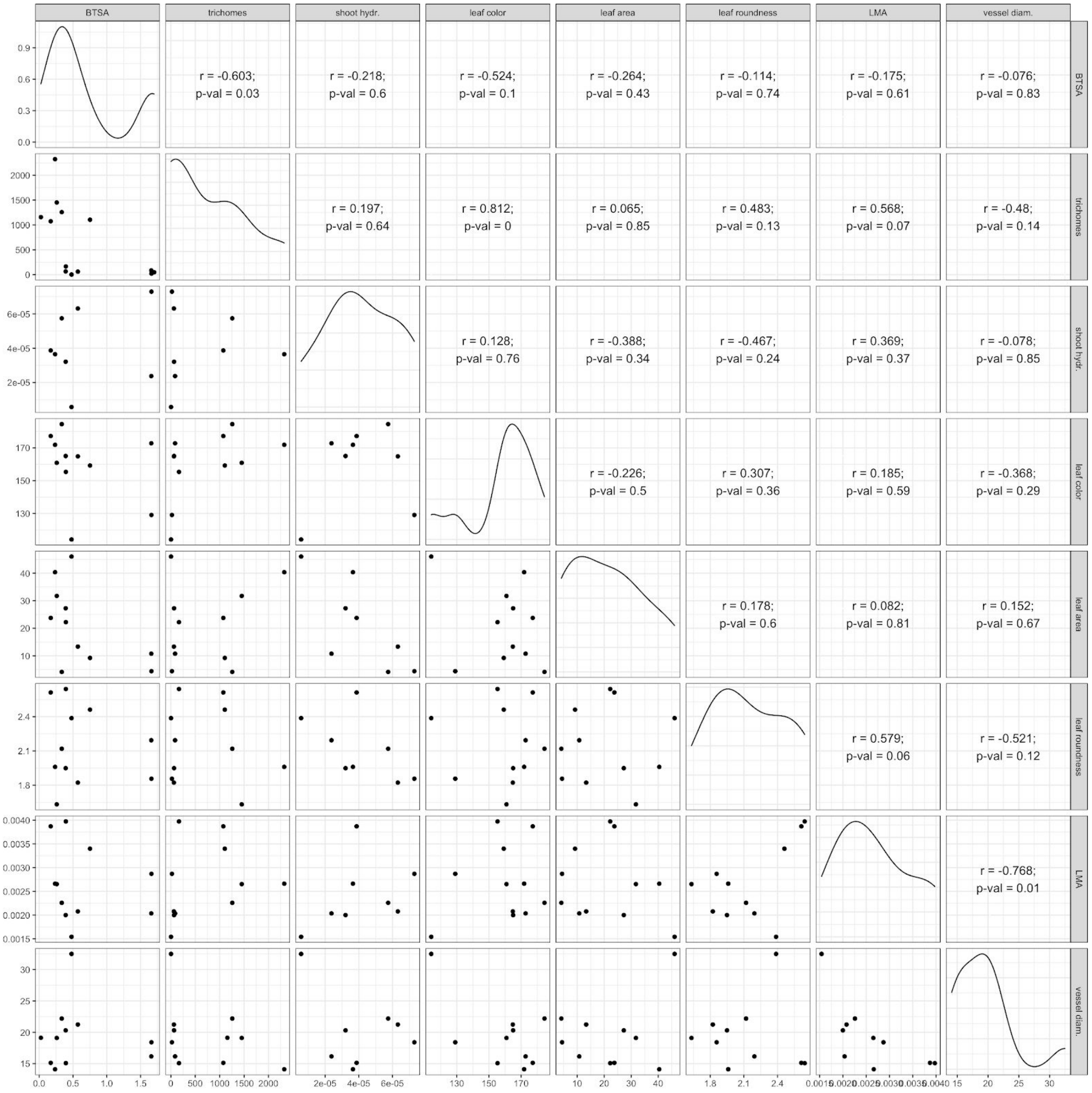


**Figure S11:** Pairwise correlations among all measured traits in *Encelia* (Table S2). Wood density was excluded due to low sample size. Shown in the upper diagonal are estimates of phylogenetic correlations and p-values between ln-transformed traits. Very few traits are correlated strongly |r > 0.6| and even fewer are significantly correlated. One of the few significant correlations is between leaf color and trichome density; this is expected as a high density of trichomes can result in a leaf looking more white.

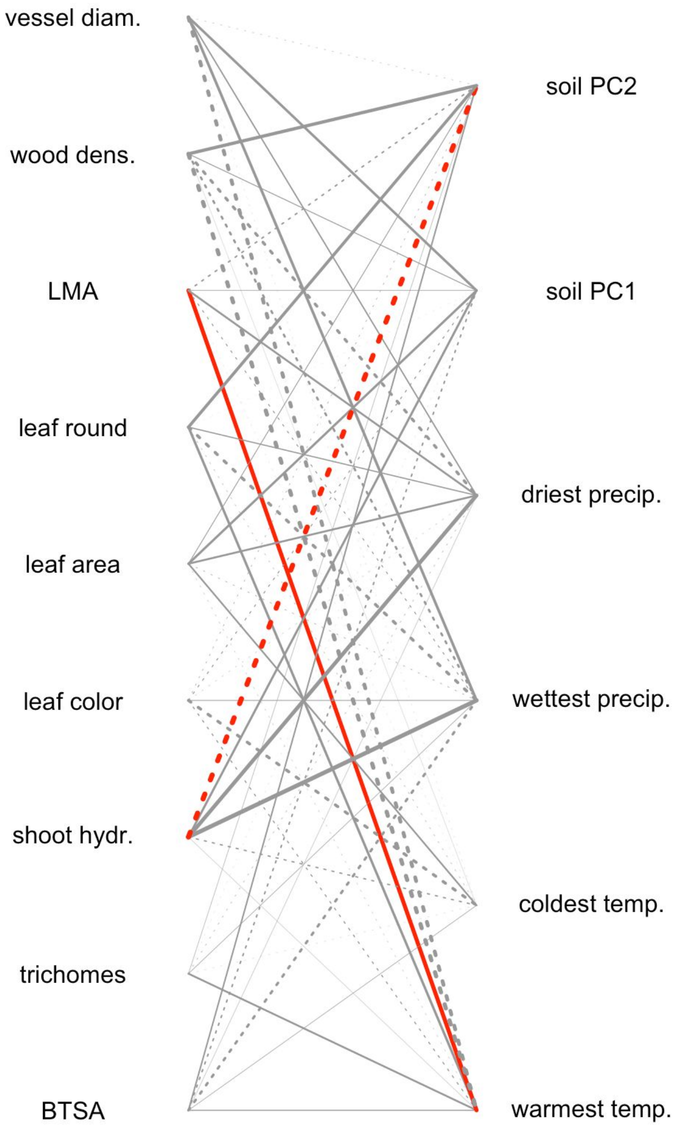


**Figure S12:** Correlations between (L) trait and (R) environmental variables. Trait abbreviations follow Table 1. Width of lines indicate absolute strength of phylogenetic correlation between ln-transformed traits and environment. Red lines indicate significant correlations ($p < 0.05$), and dotted lines indicate negative correlations. Very few traits are correlated significantly with environmental variation.
